# The umbrella cell keratin network: organization as a tile-like mesh, formation of a girded layer in response to bladder filling, and dependence on the plectin cytolinker

**DOI:** 10.1101/2024.06.11.598498

**Authors:** Wily G. Ruiz, Dennis R. Clayton, Tanmay Parakala-Jain, Marianela G. Dalghi, Jonathan Franks, Gerard Apodaca

## Abstract

The keratin cytoskeleton and associated desmosomes contribute to the mechanical stability of epithelial tissues, but their organization in bladder umbrella cells and their responses to bladder filling are poorly understood. Using super-resolution confocal microscopy, along with 3D image reconstruction and platinum replica electron microscopy, we observed that the apical keratin network of umbrella cells was organized as a dense tile-like mesh comprised of tesserae bordered on their edges by cortical actin filaments, filled with woven keratin filaments, and crosslinked by plectin. A band of keratin was also observed at the cell periphery that was linked to the junction-associated actin ring by plectin. During bladder filling, the junction-localized desmosomal necklace expanded, and a subjacent girded layer was formed that linked the keratin network to desmosomes, including those at the umbrella cell-intermediate cell interface. Disruption of plectin led to focal keratin network dissolution, loss of the junction-associated band of keratin, perturbation of tight junction continuity, and loss of cell-cell cohesion. Our studies reveal a novel tile-like organization of the umbrella cell keratin cytoskeleton that is dependent on plectin, that reorganizes in response to bladder filling, and that likely serves to maintain umbrella cell continuity in the face of mechanical distension.

## INTRODUCTION

Umbrella cells, which form the superficial layer of the stratified urothelium, must adapt to large changes in wall tension as urine is stored and then voided while maintaining an impermeable barrier ^1^. To understand how this is accomplished, one must identify the cellular structures and adaptations that make such a feat possible. Mechanisms described to date include changes in cell shape: umbrella cells become flat and squamous as the bladder fills but assume a somewhat cuboidal shape after voiding ^2, 3^. In addition, umbrella cells dramatically expand their apical surface area during filling (> 100%), a consequence of exocytosis of membrane-rich discoidal/fusiform-shaped vesicles (DFVs) ^2, 4, 5^, followed by the rapid and complete recovery of excess apical membrane by endocytosis in response to voiding ^5, 6^. Furthermore, components of the apical junctional complex (AJC), including the belt-like tight and adherens junctions and the AJC-associated actin belt undergo expansion during filling and contraction after voiding ^7, 8^. While desmosomes and keratins impart cells and tissues with mechanical strength and stability ^9–13^, and mutations in these components lead to loss of cell cohesion and tissue integrity ^9, 10, 14–17^, we have few insights into how the bladder cycle impacts umbrella cell desmosomes and the associated keratin cytoskeleton.

Within the AJC, desmosomes form a necklace of spot-like adhesions (0.2-0.5 µm in diameter) that are positioned just below the adherens junction ^18, 19^. However, desmosomes are also found along the basolateral surfaces of epithelial cells ^18^. Key structural components of the desmosome include the cadherin-family membrane proteins desmogleins (DSGs; four isoforms) and desmocollins (DSCs; three isoforms), which assemble in *cis* on their respective cell surface and then interact *in trans* with DSGs/DSCs on adjacent cells to form sites of cell-cell adhesion ^9^. Bound to the cytoplasmic tails of the DSGs/DSCs are the armadillo family proteins junction plakoglobin (JUP) and the plakophilins (PKP, isoforms 1-3), which in turn interact with the plakin family protein desmoplakin (DSP)^10^. The latter binds to keratin intermediate filaments, which are divided into type I (acidic) and type II (basic) subfamilies ^16^. Both DSG2 and DSP are load sensing ^20, 21^, indicating they are likely sites of mechanotransduction.

Keratins are well suited for bearing mechanical loads ^13, 22^. In general terms, intermediates filaments are thought to be more flexible and much less rigid than actin or microtubules ^23, 24^. In the case of keratins, they are known to straighten when stretched, they exhibit elasticity (reaching strains up to 100% and are also compressible), and they resist breakage by becoming stiffer as strain increases ^22, 25, 26^. An example of this capability is observed in situations of high tension, where an unusually straight and load bearing keratin network forms that compensates for the dilution of the actin cytoskeleton observed in MDCK cells undergoing “extreme” strains (up to 1,000%) ^27^. Keratins can also contribute to cell stiffness, and when keratin-deficient keratinocytes are probed using atomic force microscopy they exhibit a “softening” that can be overcome by expression of KRT5/KRT14 ^28^. Keratin-deficient keratinocytes also exhibit a greater propensity toward invasion ^29^. The elastic properties of keratins, along with their role in imparting mechanical resilience, would be ideal in the setting of the umbrella cell, but how or if the organization of the umbrella cell-associated keratin cytoskeleton adapts to mechanical loads remains obscure.

In epithelial cells, keratins often exhibit a loose mesh-like appearance with desmosome-associated keratins organized into functional “rim and spoke” components ^30–33^. The spokes are comprised of radial keratin filaments, which run perpendicular to the plasma membrane, linking peripheral desmosome-attached keratins to the outer nuclear envelope, likely via the LINC (linker of nucleocytoskeleton and cytoskeleton) complex ^32, 34^. Thus, the spokes serve to support the cell in response to mechanical loads and convey instances of mechanical deformation to the nucleus. The “rim” component is comprised of a thin circumferential ring of keratin filaments that runs parallel to the plasma membrane, interconnects the desmosomes, and which is hypothesized to assist in positioning and tensioning the desmosomes, particularly in response to mechanical deformations ^30, 35^. If other patterns of keratin organization exist remains an open question, but the keratin cytoskeleton at the apical pole of umbrella cells is reportedly akin to a woven mesh formed by keratins and with openings housing unfused DFVs ^36^. In addition, a “frame” of keratins is observed at the cell periphery. How individual keratin filaments are organized within the umbrella cell network, the relationship of the keratin network to AJC-associated desmosomes, and the nature of the keratin frame are unknown.

The functions and organizing principles of the rim and spoke keratins are not yet fully understood, but formation of rim-associated keratins depends on the presence of desmosomal cadherins and DSP, which recruit keratins and promotes their elongation into rim (and spoke) components ^37^. Moreover, a recent study implicates PLEC (plectin) in their formation ^35^. PLEC is a large (>500kD) ubiquitously expressed cytolinker that binds actin, keratin, and microtubules, recruiting them to a variety of organelles including mitochondria, the nucleus, and hemidesmosomes ^38, 39^. In cultured MDCK cells, knockout of *Plec* expression, or the use of a selective PLEC inhibitor called plecstatin-1 ^40^, reveals roles for PLEC in mediating keratin-actin interactions at the AJC, formation of the keratin rim, organization of the spokes, planar arrangement of desmosomes, and overall monolayer cohesion ^35^. PLEC may also contribute to cell-cell adhesion, as conditional hepatocyte/cholangiocyte KO mice and conditional intestinal epithelial KO mice exhibit junctional defects ^41, 42^. If PLEC plays a similar role in organizing the apical keratin network of native epithelia such as the urothelium remains to be described, although several epithelial tissues are known to express this protein ^43^.

Our studies reveal that instead of a loose mesh with rim and spoke components, the keratin cytoskeleton of native umbrella cells is organized as a dense apical tile-like mesh comprised of bordering actin filaments, filled with woven keratin filaments, and crosslinked by cytolinkers. Furthermore, a thin band of keratin is observed at the cell periphery, which is linked to the AJC-associated actin belt by PLEC. During bladder filling we observe the formation of a basal girded layer that functions to link the keratin network to desmosomes including those at the umbrella cell-intermediate cell junction. The continuity of the umbrella cell keratin network depends on PLEC and disruption of this cytolinker leads to focal keratin network dissolution, defects at the AJC, and loss of cell-cell cohesion. Our observations support the likelihood that the keratin cytoskeleton and desmosomes contribute to umbrella cell cohesion during the bladder filling/voiding cycle.

## RESULTS

### AJC-associated desmosomes, comprised of DSG2 and DSC2, form a necklace-like structure that expands/contracts with bladder filling and voiding

To understand how filling and voiding impact umbrella cell desmosomes, we initially sought to define desmosome morphology, organization, and protein composition. When examined using thin-section transmission electron microscopy (TEM), the AJC of the rat bladder umbrella cells was comprised of readily identifiable tight junctions, adherens junctions, and desmosomes (Fig. 1A). The latter were positioned just below the adherens junction, where a characteristic ∼ 25nm gap, filled with electron-dense material, formed between adjacent cells ^18^. A dense cytoplasmic plaque was also noted on either side of the junction. A mass of cytoskeletal filaments (labeled “Csk” in Fig. 1A) was observed adjacent to the desmosomes, but these elements were abundant along the entire aspect of the AJC. Single-cell transcriptomic data obtained from closely related mouse umbrella cells indicates that *Dsg2* and *Dsc2* are likely the sole or major desmosomal cadherins expressed by these cells ^44^. We confirmed that DSG2 and DSC2 were co-expressed in spot-like structures at the interface of adjacent rat umbrella cells, both at the AJC and along the lateral membranes (Fig. 1B).

**Fig. 1.**
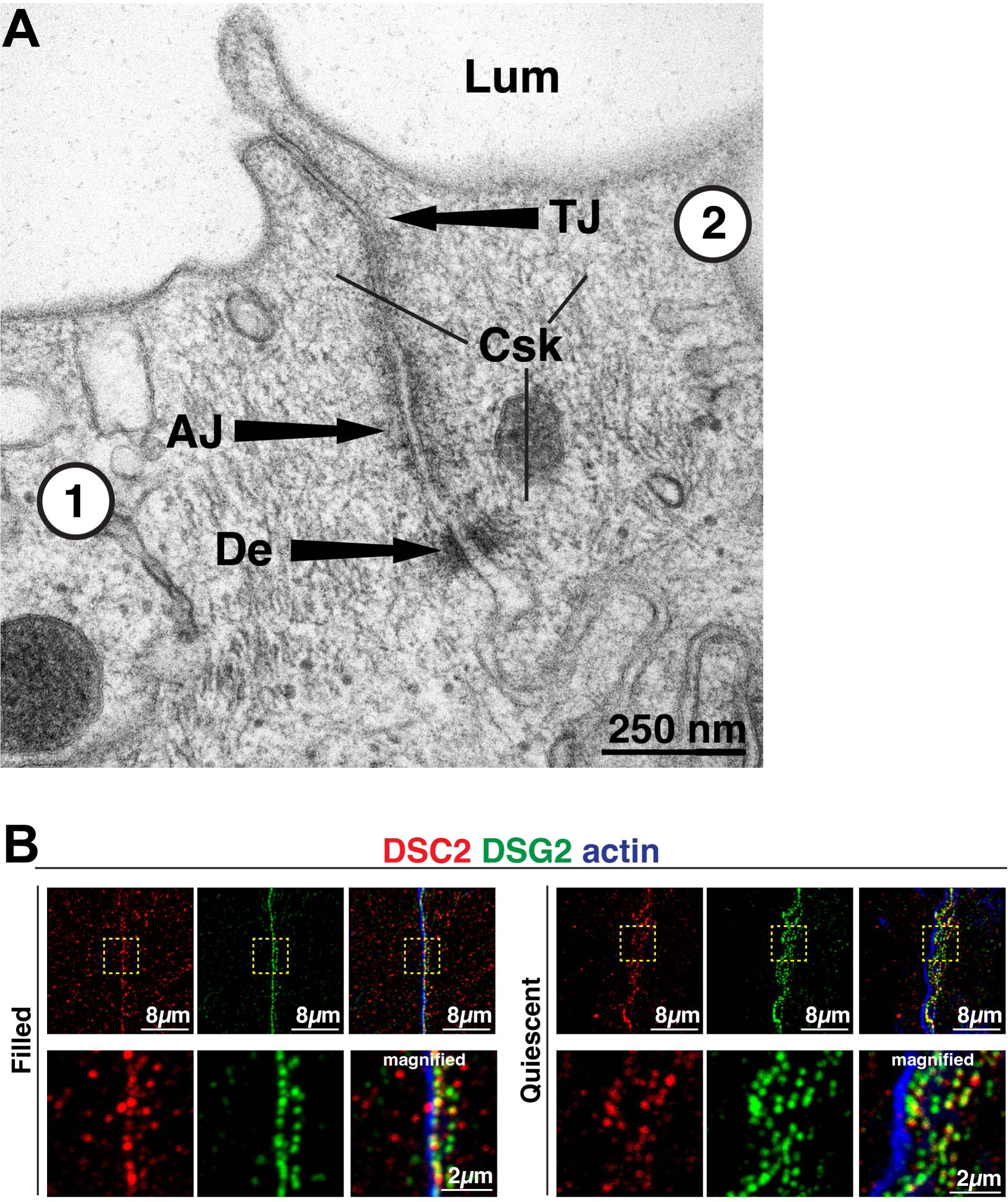
Bladder umbrella cell desmosomes. **(A)** Thin-section transmission electron micrograph of rat bladder umbrella cells. An apical junctional complex (AJC)-associated desmosome is localized below the adherens junction (AJ) and tight junction (TJ) of two adjacent cells (labeled 1 and 2). Legend: Csk, cytoskeletal filaments; De, desmosome; Lum, lumen. **(B)** Expression of DSC2 and DSG2 at the AJC of two adjacent umbrella cells. Phalloidin was used to label the AJC-associated actin ring, which is positioned above desmosomes. Extending to the right of the AJC-associated desmosomes are those found along the lateral surfaces of the adjoining cells. Images are 3D reconstructions of confocal Z stacks. The boxed regions in the upper panels are magnified in the panels below.

To better understand the organization of the desmosomes in the umbrella cell AJC, we performed confocal microscopy, coupled with super resolution image processing (SRIP; nominal resolution of 120 nm), and 3D surface rendering (see Video 1). This analysis revealed that the desmosomes could be discriminated from the apical-most tight junction (labeled with TJP1, aka ZO1) and from the belt of actin that lay between TJP1 and the adherens junction (Fig. 2A-B). The latter is associated with the upper 1/3 of CDH1 staining, whereas the lower 2/3 of CDH1 staining was associated with the lateral membranes of the umbrella cell and overlapped with DSG2. Given the small gap between apposing membranes in desmosomes, it was not possible to resolve the desmosome into two halves. Instead, desmosomes had a beaded appearance and were situated at the interface between adjacent umbrella cells. We identified three populations of desmosomes in umbrella cells. The first of these were AJC-associated (magenta-colored in Fig. 2). These were found just below the adherens junction and are equivalent to the AJC-associated desmosomes identified by TEM above (Fig. 1A). These desmosomes had an ovoid appearance, and formed a necklace-like structure that surrounded the apex of the cell. However, at the tricellular region of the AJC, which characteristically exhibits a dip in junction height (see arrowheads in Fig. 2A)^45^, DSG2 was absent from the base of the structure (see also inset in Fig. 3A, upper panel). In contrast, TJP1, actin, and CDH1 formed continuous belts. The second population of desmosomes, those along the lateral surfaces of the umbrella cell, were sparser and did not form a necklace-like structure (cyan-colored in Fig. 2B-C). The third population were found at the base of the umbrella cell forming sites of cell-cell contact between umbrella cells and the underlying intermediate cells (blue-colored in Fig. 2B-C).

**Fig. 2.**
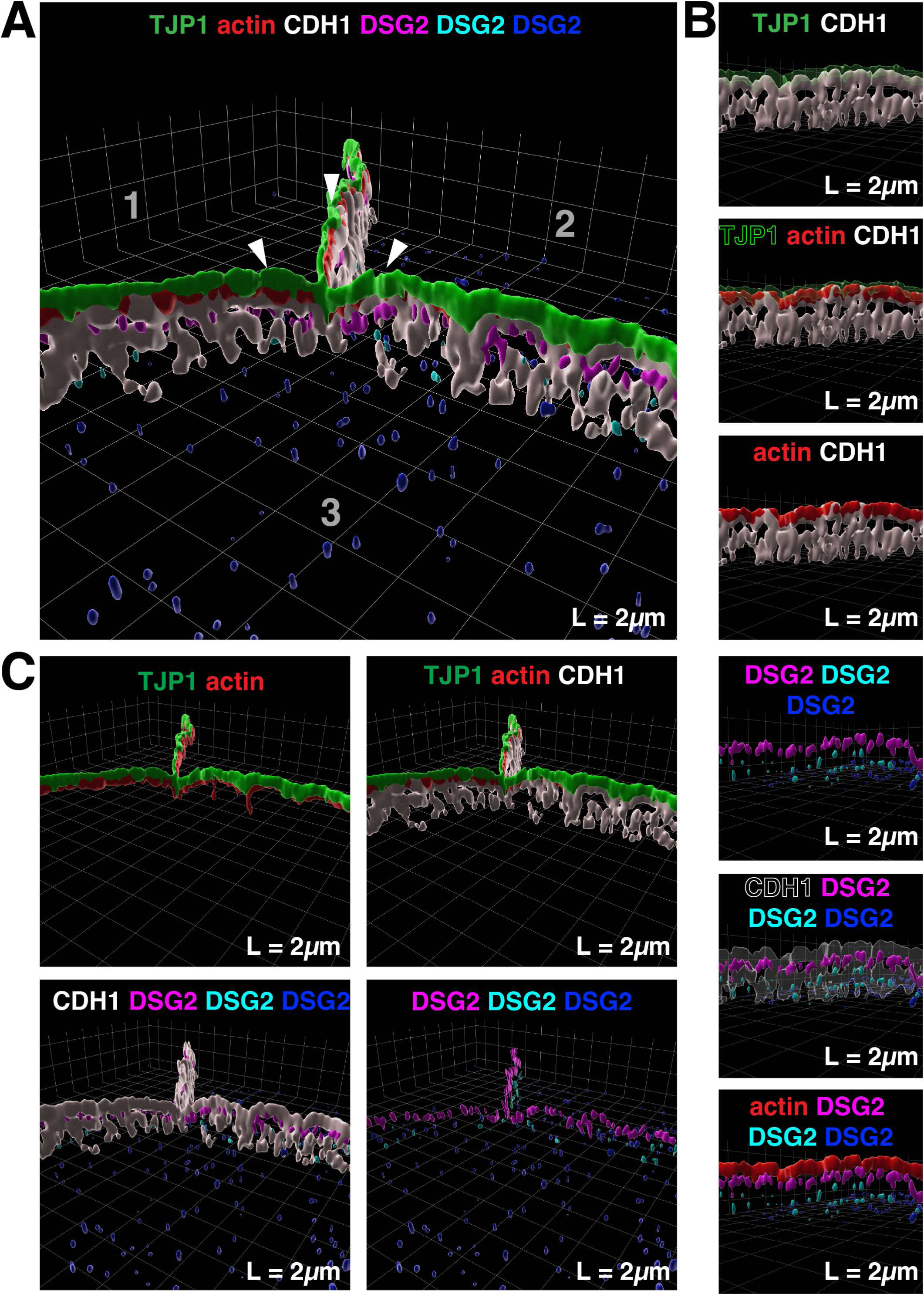
3D reconstruction of the umbrella cell AJC. **(A)** 3D surface reconstruction of a confocal-SRIP Z-stack taken at the tricellular region of three adjacent umbrella cells (labeled 1-3) obtained from a filled bladder. Arrowheads indicate the site where the tricellular junction dips. **(B)** The larger image was rotated along its lateral axis and digitally sliced to allow a head-on view of the AJC in the bicellular region between two cells. **(C)** Additional views of tight junction, adherens junction, AJC-associated actin belt, and desmosomes. DSG2-labeled desmosomes were classified as being AJC associated (magenta), lateral surface associated (cyan), or found at the interface of umbrella cells and subjacent intermediate cells (dark blue). **Video:** See companion Video1 for 3D rendering of AJC.

**Fig. 3.**
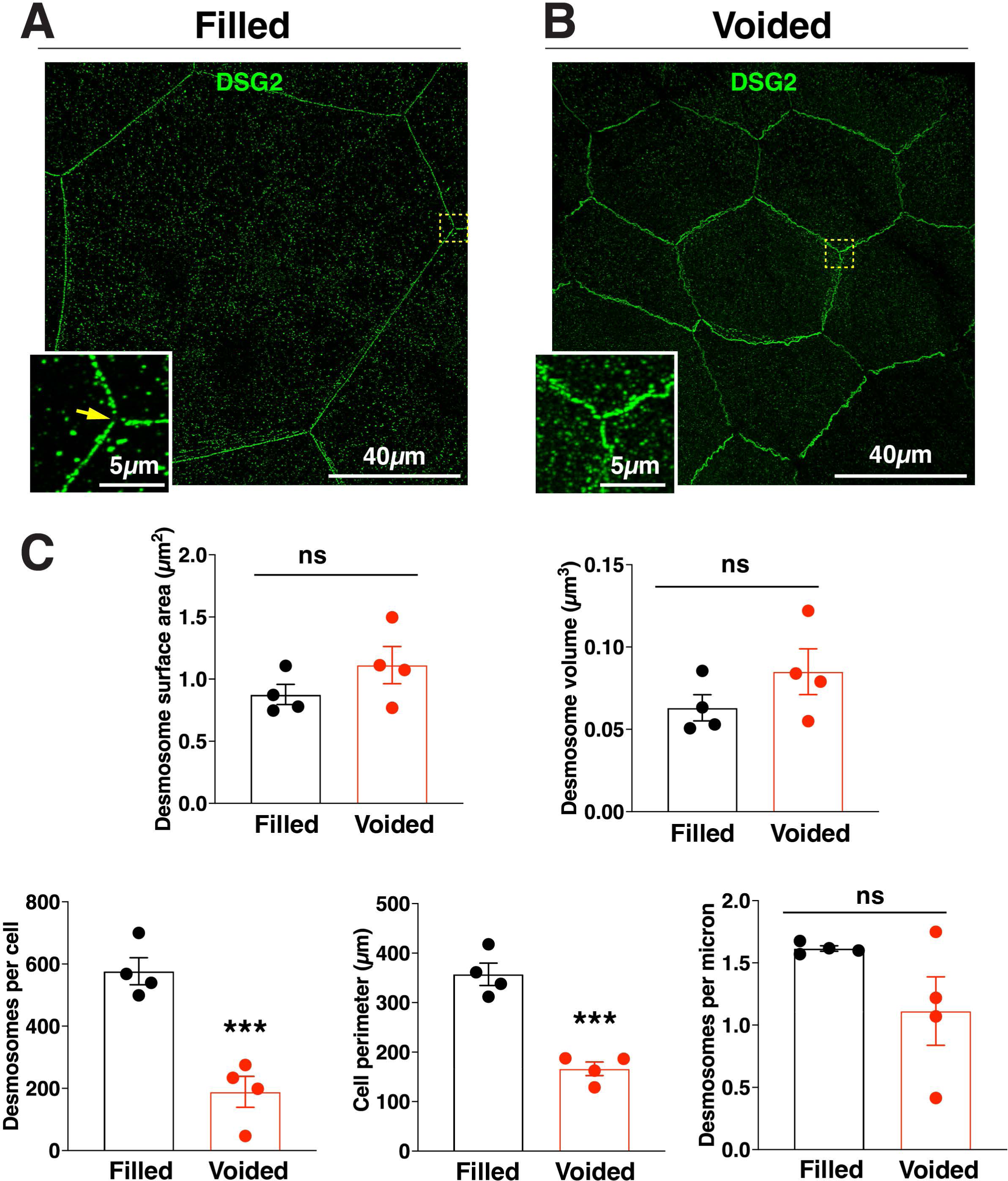
Effects of bladder filling and voiding on the AJC-associated desmosome necklace. **(A-B)** Representative examples of DSG2-labeled umbrella cells taken from filled (A) or voided (B) bladders imaged using confocal-SRIP microscopy and rendered in 3D. The boxed regions are magnified in the insets. The yellow arrowhead in the inset of panel A indicates the site of the tricellular junction. **(C)** Quantification of the indicated parameters for filled and voided bladder umbrella cells. Data are mean ± SEM (n=4 animals). Significant differences are indicated by asterisks, and non-significant ones indicated by “ns.”

Like the tight junction and adherens junction of umbrella cells ^7^, we observed that the AJC-associated necklace of desmosomes expanded during filling and shrank after voiding (Fig. 3A). Absent of the waviness of the cell borders in voided bladders, the continuity and organization of AJC-associated desmosomes were not obviously affected by these events (Fig. 3A-B). In either case, we observed a continuous necklace of desmosomes surrounding the cell periphery (absent of regions at the base of tricellular junctions). Furthermore, and at this level of resolution, there was no significant difference in average desmosome surface area or average desmosome volume between the two sample populations (Fig. 3C). In contrast, the average number of desmosomes per cell was significantly greater in umbrella cells taken from filled bladders (∼600) vs voided bladders (∼200)(Fig. 3C). This was concordant with the greater cell perimeter of filled vs voided umbrella cells (Fig. 3C). Interestingly, when the number of desmosomes per µm of perimeter length was calculated, there was no significant difference between filled and voided bladders (Fig. 3C). This may indicate a cellular mechanism that holds the ratio of desmosomes/cell perimeter length relatively constant even in the face of large changes in cell perimeter as the bladder fills and empties. In sum, like the tight and adherens junction, the umbrella cell’s necklace of desmosomes also accommodates the bladder’s filling and voiding cycle.

### The umbrella cell keratin cytoskeleton is organized as an apical mesh and a subjacent girded layer that forms during bladder filling

To further explore the organization of the umbrella cell keratin network, we localized KRT20 in cross sections of urothelium derived from filled and quiescent bladders (the latter were emptied, but not allowed to fill). Umbrella cells from filled bladders exhibited a squamous morphology, while the umbrella cells derived from quiescent bladders had a roughly cuboidal shape, often with a basal pedicle housing the cell’s two nuclei (Fig. 4A). In both cases, KRT20 amassed at the apical pole of umbrella cells forming a relatively thick mesh-like structure, while less KRT20 was found at the basal pole of the cells (Fig. 4A). The underlying intermediate and basal cells were KRT20 negative.

**Fig. 4.**
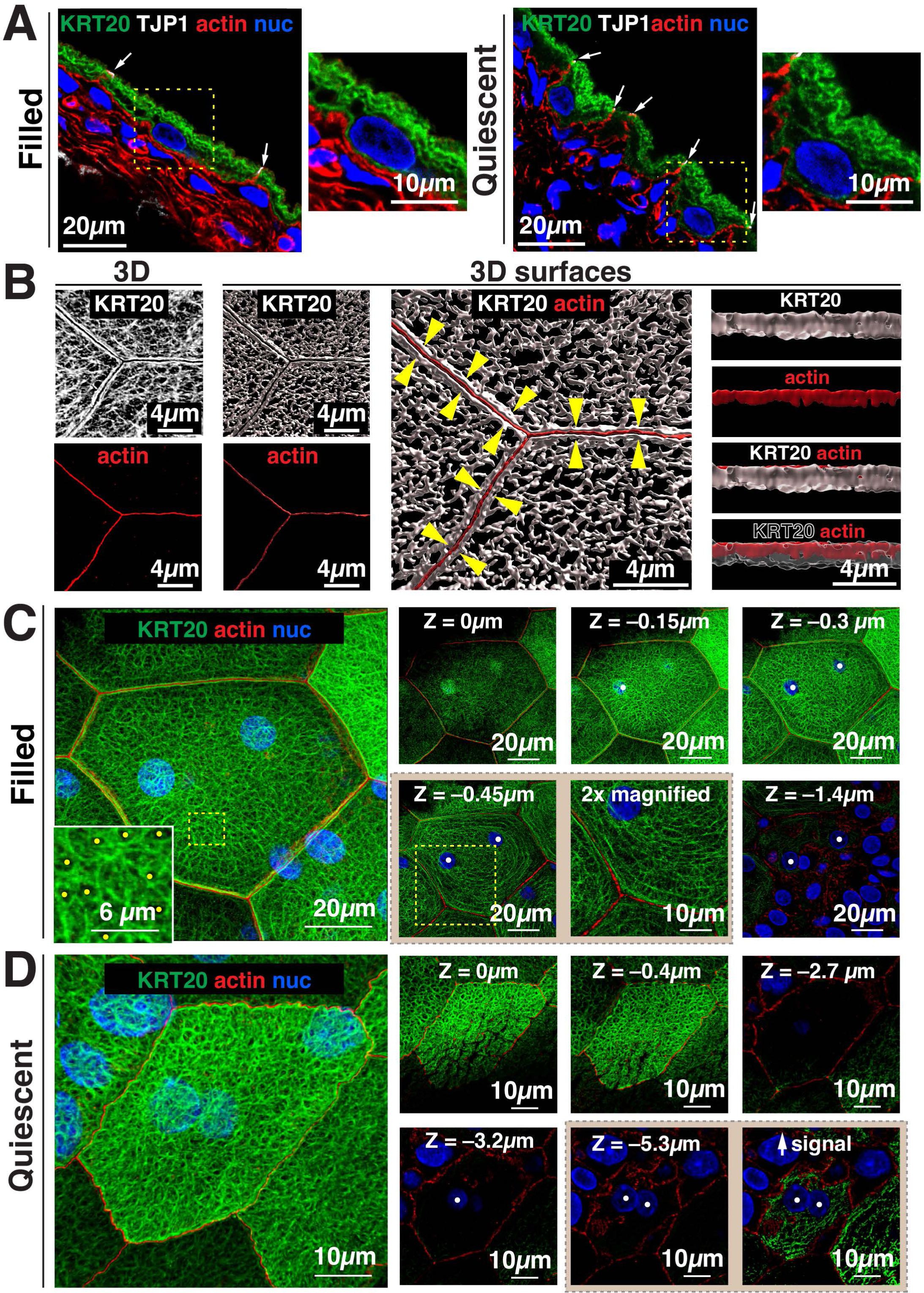
Organization of the keratin network in umbrella cells obtained from filled and quiescent bladders. **(A)** Cryosections of urothelium taken from filled or quiescent bladders. White arrows indicate the site of the AJC, and boxed regions are magnified in the smaller panels to the right of the larger images. **(B)** Reconstruction of the umbrella cell KRT20-labeled network in 3D or as 3D surfaces (derived from confocal-SRIP images obtained from filled bladders). Yellow arrowheads indicate the band of keratin at the cell periphery. Panels to the right: the adjacent overview image to the left was rotated 90°along its lateral axis and digitally sliced allowing for a head-on view of just the region of the keratin band near the AJC. **(C-D)** Organization of the KRT20-labeled network in an umbrella cell taken from a filled bladder (C) or a quiescent bladder (D) and imaged using confocal-SRIP microscopy. **(C)** The boxed region in the image to the left is magnified in the inset. Yellow dots indicate locations of lacunae. Images to the right are individual X-Y-sections taken at the indicated depth (apical-most section was set to Z=0). The position of nuclei is indicated by white dots. Boxed region in Z = -0.45 µm is magnified 2-fold in the image to the left. **(D)** The KRT20 signal in Z = -5.3 µm was increased in the panel to the right. **Videos:** Companion videos show the apical-most aspect of the keratin network of umbrella cells taken from filled bladder (Video2) or the girded layer (Video3). A video is also presented for an umbrella cell taken from a quiescent bladder (Video4).

The complexity of the KRT20 network became apparent when whole-mounted urothelium, obtained from bladders filled to one-half their normal capacity (∼ 0.5ml for rats of this weight), was imaged *en face* using confocal-SRIP microscopy and 3D surface reconstruction (see Video 2). At the periphery of each umbrella cell, we observed a thin continuous band of KRT20 that collected on either side of the AJC-associated actin ring (see yellow arrowheads in middle right panel of Fig. 4B). This thin peripheral band of KRT20 corresponds to the “frame” described previously by Veranic and Jezernik ^36^. Analysis of 3D surfaces revealed that the peripheral band of KRT20 extended up to the level of the AJC-associated actin ring (right most panels, Fig. 4B), although the KRT20 band was sometimes shorter. Away from the AJC, we observed that the keratin network was polarized along the apical-basal axis, forming two somewhat overlapping layers, which were best visualized in individual X-Y sections taken from confocal Z-series. An upper apical “mesh-like layer” was composed of keratin filaments that appeared as interlaced strands sometimes forming lacunae (i.e., small spaces or gaps) with diameters in the range of 0.4-0.8 µm (see inset in left-most panel of Fig. 4C). These lacunae were previously shown to house clusters of DFVs prior to their fusion ^36^. The adjacent areas of the mesh were more tightly woven and lacked lacunae. In umbrella cells taken from full bladders, two circular-shaped bumps of keratin were often observed in the upper portion of the mesh layer, which were positioned directly above the paired nuclei of these cells (see optical section Z = 0 in Fig. 4C).

Below the mesh layer of these filled bladders, and extending deeper into the cytoplasm, was a basal “girded layer” comprised of keratin filaments that in X-Y sections were organized in a belt-like pattern of nested ovals that followed the contours of the cell perimeter (see optical sections Z = –0.3 and Z = –0.45 in Fig. 4C). The ovals below the mesh layer were cross-linked by struts comprised of perpendicularly arranged keratin filaments (see 2X magnified panel in Fig. 4C)(see also Video 3). In these filled bladder preparations, the upper aspects of each of the umbrella cell’s two nuclei were embedded within these nested ovals, apparently connecting the nuclear ectocytoskeleton to the keratin network. However, there was relatively less keratin signal associated with the basal pole of these nuclei (Z = –1.4 in Fig. 4C). Given that keratin filaments are comprised of heteromers of acidic and basic keratins ^16^, it was not surprising that KRT8 (a basic keratin) and KRT20 (an acidic keratin) shared an overlapping distribution in umbrella cells (Supplemental Figure S1); however, KRT8 was also expressed by the underlying intermediate and basal cell layers. We also assessed the organization of KRT20 in umbrella cells in which the bladder was filled to capacity (1.0ml). The umbrella cell layer remained patent, the cells were very flat, they exhibited maximal diameters, and their keratin network appeared similar to what is described above (Supplemental Figure S2). However, the girded layer was more pronounced in these cells, possibly indicating more recruitment of keratin to this layer to support the increased wall tension.

We also examined the organization of the keratin network in the umbrella cells of quiescent bladders. In these preparations, the umbrella cells exhibited apical folding (apparent as discontinuities of the keratin network when individual optical sections are examined, e.g., see Z = 0 and Z = –0.4 µm, Fig. 4D), and like filled bladders, the apical keratin network had a mesh-like appearance (Fig. 4D and Video 4). Strikingly, there was an absence of a visible girded layer in quiescent umbrella cells and relatively little KRT20 signal was associated with the nuclei of these cells (Fig. 4D and Video 4). However, KRT20 could be detected if the signal intensity was boosted (see right-most panel, Z = –5.3µm, Fig. 4D). In sum, the apical keratin network in umbrella cells is comprised of a mesh-like layer and a subjacent girded layer. The latter is not observed in quiescent bladders but develops in response to bladder filling and is presumably a mechanism to maintain tissue cohesion in the face of increased wall tension.

### The umbrella cell KRT20 network interacts with AJC-associated and basal desmosomes

We next sought to understand the relationship of the keratin network and the desmosomes, important sites of keratin:membrane attachment in other cell types ^10^. Foci of the keratin-binding protein DSP were localized to either side of DSG2-labeled desmosomes, giving the impression of parallel, registered, and dashed lines of DSP when adjacent cells were viewed from above (Fig. 5A). 3D surface reconstructions revealed that DSP and DSG2 were roughly within the same Z-plane and the tripartite structure was oriented parallel to the cell surface (see Fig. 5A-B). To better understand the relationship of KRT20 to DSP, 3D surface reconstructions were generated. They revealed that between adjacent cells the peripheral band of KRT20 formed a fjord-like structure with DSP-DSG2 at its base (see Video 5). XZ slicing further confirmed that the peripheral band of KRT20 (along with most of the apical mesh layer) extended above the DSP/DSG2-labeled desmosomes (Fig. 5B); although, the lower quarter of the KRT20 band overlapped with DSP (Fig. 5B), which likely serve as potential sites of KRT20 anchorage.

**Fig. 5.**
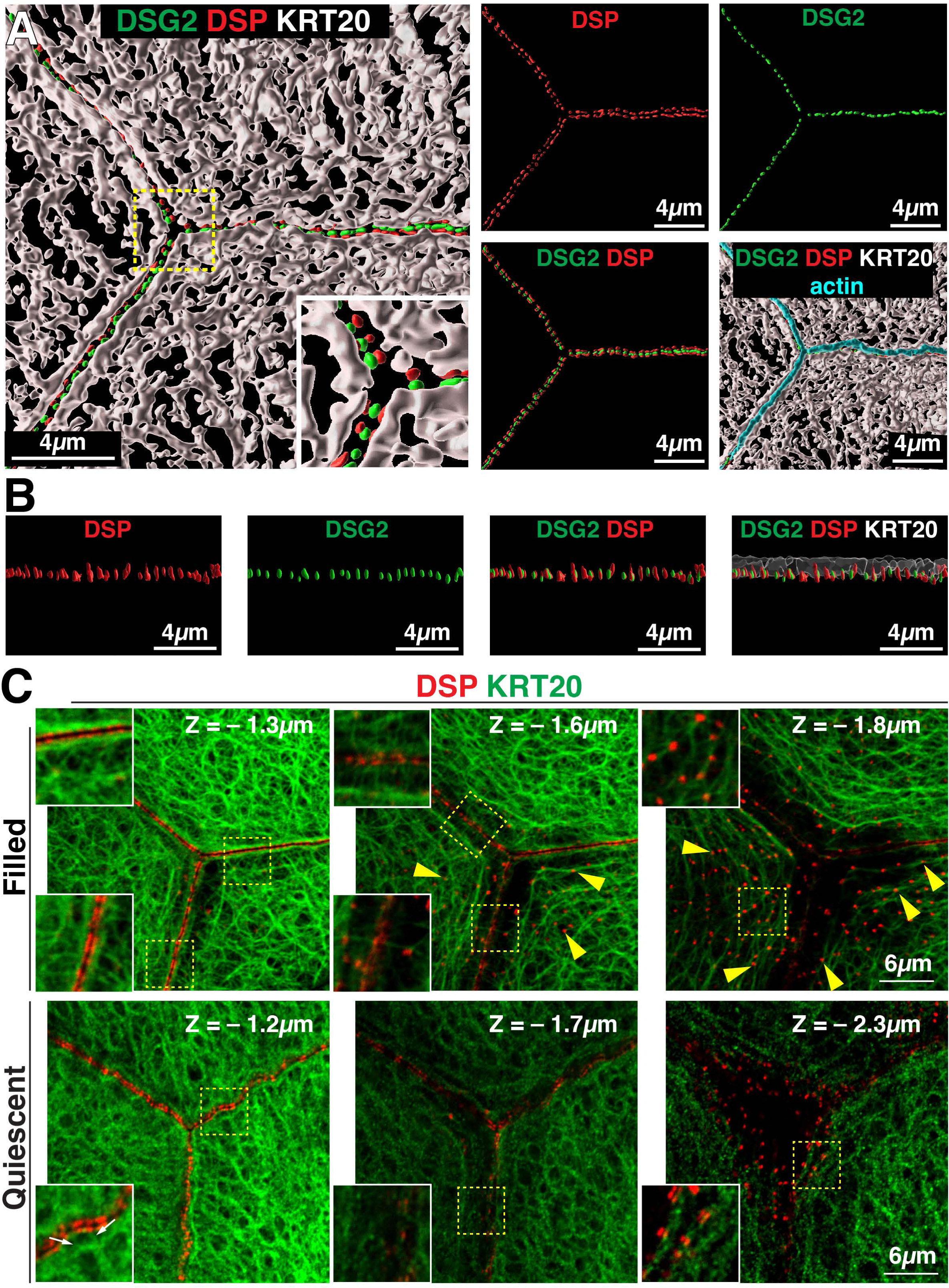
Localization of DSG2, DSP, and KRT20 in umbrella cells of filled and quiescent bladders. **(A)** 3D surface reconstruction of confocal-SRIP Z-stack showing the DSG2, DSP, KRT20, and the AJC-associated actin ring of umbrella cells taken from filled bladders. **(B)** The upper left image was rotated 90°along its lateral axis and digitally sliced allowing for a head-on view of just the region of the AJC-associated desmosomes and KRT20 network. **(C)** A tricellular region of the umbrella cell layer, taken from filled or quiescent bladders, was imaged using confocal-SRIP microscopy. X-Y sections at the indicated Z level are shown (apical-most section is Z = 0). Boxed regions are magnified in the insets. Yellow arrowheads indicate examples of close association between the girded layer and desmosomes at the umbrella cell/intermediate cell junction. Small white arrows in lower left panel indicate short keratin struts in quiescent samples connecting the desmosomes and the adjacent KRT20 mesh. **Video:** A companion video shows the 3D relationship between the apical keratin network and the DSP/DSG2-labeled desmosomes (Video 5).

To better understand the sites of KRT20/desmosome interaction, we scrutinized X-Y optical sections taken from the tricellular region of filled bladders. In Z planes with the brightest DSP signal, we observed that the basal aspects of the peripheral KRT20 band came near the AJC-associated DSP signal (left-most upper panel in Fig. 5C). In addition, KRT20 struts emanated from either side of the junction that linked the AJC-associated keratin to the remaining mesh-like network. The straight appearance of these struts is consistent with keratin filaments under tension ^22^. In the girded layer, the nested oval KRT20 filaments appeared to be organized by contact with desmosomes at the umbrella cell/intermediate cell junction (middle and left-most upper panels of Fig. 5C). In quiescent bladders, we observed that the KRT20 band near the AJC-associated desmosomes was offset a small amount from the DSP signal by what appeared to be short struts or cross bridges that connected to filaments that ran parallel the membrane (see inset in bottom right-most panel of Fig. 5C). At the base of the quiescent umbrella cells, a small fraction of basal desmosomes appeared to associate with KRT20 filaments (lower middle panel of Fig. 5C).

In sum, the keratin network of umbrella cells taken from filled bladders interacted with desmosomes at two sites: (i) those within the AJC and (ii) those at the base of the umbrella cells. At the AJC, the interaction was of a rim-and-strut type that connected the keratin network to the mesh layer, whereas at the base of the cells the girded layer of keratin appeared to be organized by interactions with desmosomes at the umbrella cell:intermediate cell interface. In quiescent umbrella cells, keratin interactions with the AJC-associated desmosomes were observed, while interactions with basal desmosomes was less pronounced.

### Electron microscopy reveals that the apical mesh-like network of umbrella cells is comprised of keratins, actin, and cytolinkers arranged in a tile-like pattern

We used electron microscopy to explore the apical cytoskeleton of umbrella cells (which were subjected to membrane extraction and cytoskeletal stabilization). In SEM micrographs, individual umbrella cells were identified by the presence of the AJC, which encircled the cell and appeared as a raised ridge at the cell periphery (Fig. 6A-B). At higher magnifications, a closely packed mesh of filamentous cytoskeletal elements became apparent (Fig. 6B-D). The filaments had a diameter of ∼ 20nm, although the nominal deposition of ∼ 2.5-4.0nm of gold-palladium would place these in the range of 12-15nm, indicating many were intermediate filaments. However, it is not possible to reliably distinguish keratins from actin (or microtubules) in these images. Most filaments appeared straight, extending for 50-400nm, but some were curved. Thinner filaments with a nominal diameter of ∼11nm (actual diameter 3-6nm) were also observed, as were thicker cables ∼30-55nm (actual diameter 22-50nm). The latter were sometimes hydra like, branching into smaller diameter fibers.

**Fig. 6.**
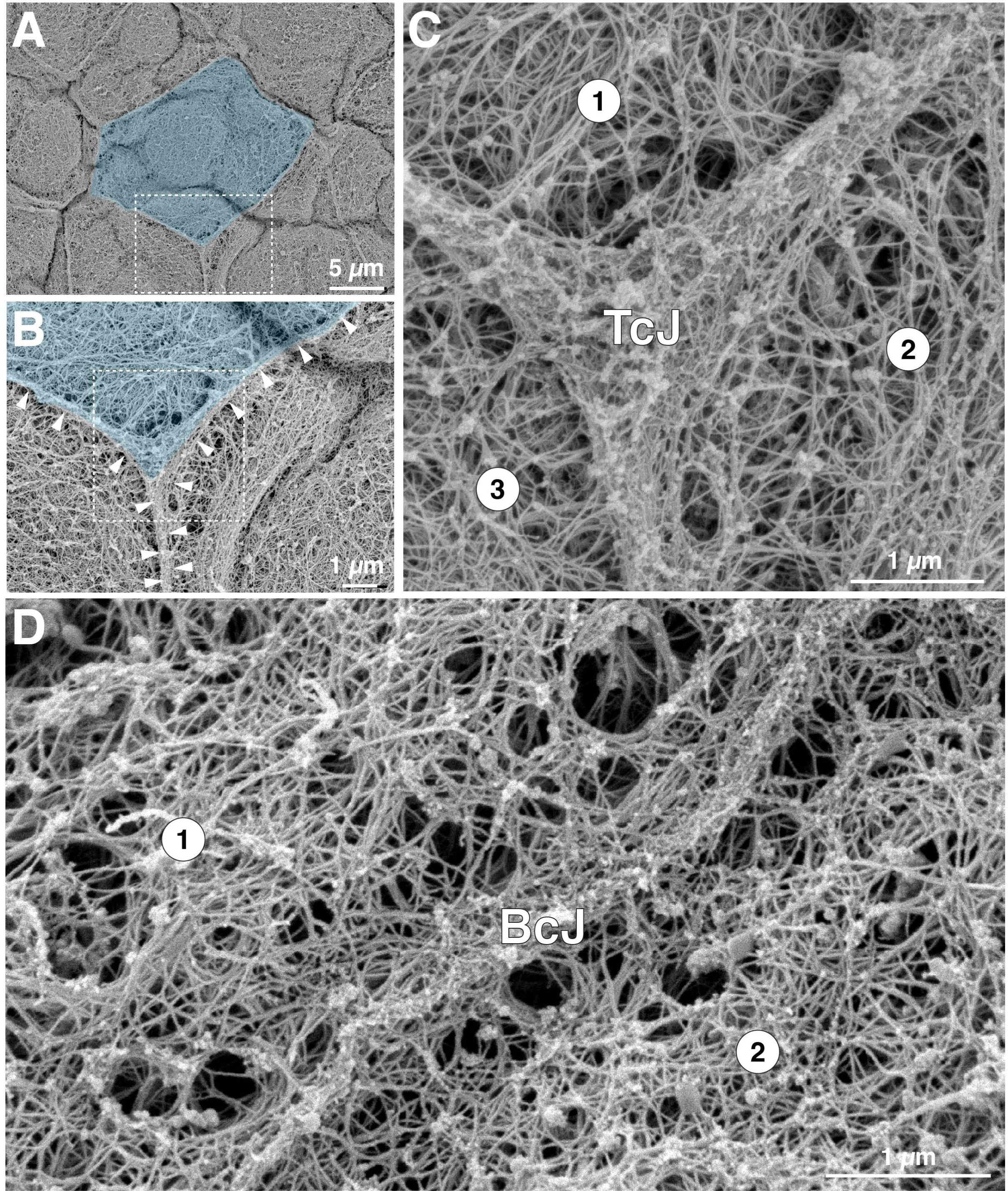
SEM analysis of the cytoskeletal network near the AJC of umbrella cells. **(A)** Overview of umbrella cell cytoskeleton revealed by detergent extraction, cytoskeleton stabilization, and SEM. An individual umbrella cell is false-colored blue. The boxed region is magnified in panel B. **(B-C)** The region of the tricellular junction (TcJ) is magnified. White arrowheads in panel B indicate the position of cell borders. The boxed region in B is magnified in panel C. The circled numbers indicate the three cells that join to form this region of the umbrella cell layer. **(D)** Organization of cytoskeleton near the bicellular junction (BcJ) of two cells labeled 1 and 2.

The microarchitecture of the filament network varied to some degree by cellular location. At the region of the tricellular junction, a dense network of crisscrossed (woven) filaments was observed to form an inverse Reuleaux triangle (three-sided polygon; Fig. 6C). Reflecting the very close apposition of membranes and cytoskeletal elements at the tight junction, it was not possible to make out cell borders in this or other regions of the AJC. At the periphery of the tricellular junction, the ∼ 20nm filaments were oriented parallel to the junction, but others appeared to extend away from the junction in a perpendicular fashion. The latter ones likely attached the AJC to adjacent filaments in the network (Fig. 6C). The bicellular regions of the AJC shared many of the features of the tricellular AJC including the lack of identifiable cell borders and a dense network of filaments that was associated with the bicellular regions of the AJC. Some of these keratin filaments ran parallel to the junction while others extended away from the junction in a perpendicular fashion (Fig. 6D). Again, the latter attached the bicellular junction to the main network.

Away from the AJC, the subapical cytoskeletal mesh appeared to be organized somewhat akin to a mosaic comprised of tesserae (tile-like structures) encased by bordering elements and filled with ∼ 20nm filaments (Fig. 7A). The size of the resulting tesserae varied: small ones had areas of ∼ 1.0µm^2^, but larger ones had areas approaching several microns (5-8µm^2^). The bordering elements had lengths on the order of ∼ 0.5-2.0µm (but sometimes extending 5 or more microns) and diameters of ∼ 75-200nm (examples of bordering elements are demarcated by white arrowheads in Fig. 7A). The constituents of bordering elements were sometimes difficult to assess as these elements were most likely to retain proteins during the extraction process; however, in regions devoid of attached proteins a filamentous substructure was apparent (and as will be noted below they contain actin). The ∼20nm diameter filaments that filled each tessera appeared to span the bordering elements, forming an elaborate crisscross pattern, or in some cases generating a central hub with emanating spokes (see white arrows in Fig. 7A). The other structure we observed in the mesh were lacunae, which had walls comprised of crisscrossing fibers that encapsulated the opening (Fig. 7B-C). A small number of filaments was observed to extend across the lacunal opening and are discussed further below (yellow arrows in Fig. 7C). Lacunae were present across the apical region of the mesh but were often found along the AJC (Fig. 7B), indicating that the AJC may be a site of active exocytosis.

**Fig. 7.**
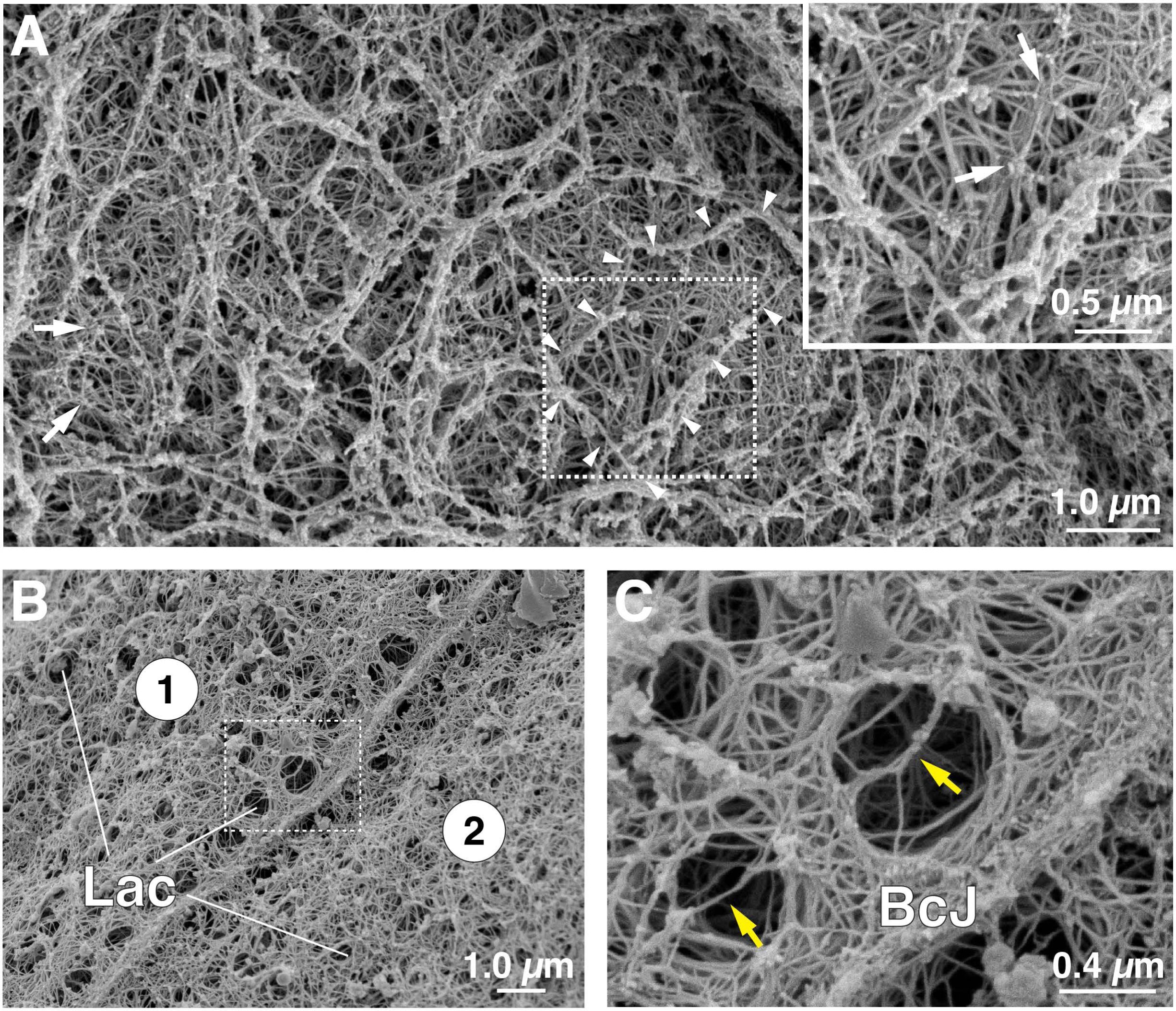
SEM analysis of umbrella cell apical cytoskeletal network. **(A-C)** SEM images taken from regions of umbrella cell apical cytoskeleton lacking lacunae (A) or those containing lacunae (B-C). **(A)** White arrowheads outline a tessera encased by thicker bordering elements. White arrows indicate regions of mesh where 20nm filaments are organized as a central hub with emanating spokes. Boxed region is magnified in inset. **(B-C)** Region near a bi-cellular junction (BcJ) where lacunae (Lac) are present. Boxed region is magnified in panel C. In C, yellow arrows indicate sparse cytoskeletal elements that traverse the lacunae.

To gain more insights into filament identity and organization, we turned to PREM, a technique usually reserved for coverslip-grown cells ^46^; however, we found that it could be performed on native urothelial tissue. In addition to aiding our recognition of keratin filaments, it also allowed us to identify short, small-diameter cytolinkers (e.g., PLEC), and to distinguish the actin cytoskeleton, which we decorated with myosin heavy chain S1 fragments ^47^. We first examined the region near the AJC, which in PREM appeared flatter than by SEM, but could be identified by its continuous belt-like architecture, its width of 200-400nm, and its granular appearance when viewed from above (it is color-coded blue in Fig. 8A). Short, unbranched actin filaments (typically < 1.0µm in length; color-coded red in Fig. 8A) with random polarity were observed to insert into the bicellular AJC and in some cases appeared to ride above it. Actin filaments were also observed in the areas away from the AJC. Keratin filaments were identified by their nominal diameter of ∼ 16nm (actual diameter of ∼12nm accounting for the ∼ 2nm platinum coating) and are color-coded green in Fig. 8A. At high magnifications, they had the appearance of a dense pipe cleaner. Keratin filaments were observed to insert into the AJC at acute angles or they paralleled the long axis of the AJC. Away from the AJC, the keratins appeared crisscrossed. The other notable structures at the AJC were small-diameter fibers (∼ 6 nm nominal diameter, ∼ 2nm actual diameter), which we tentatively identified as cytolinkers (orange-colored structures in Fig. 8A) that primarily crosslinked the intermediate filaments to one another. These appear similar if not identical to PLEC cytolinkers previously described and that have a diameter of ∼ 2-3nm ^48, 49^. We also observed cytolinkers that extended from the AJC to adjacent filaments. At the region of the tricellular junction, we again noted short, randomly oriented actin filaments accumulating near the junction, as well as the presence of keratin filaments and short cytolinkers (Fig. 8B).

**Fig. 8.**
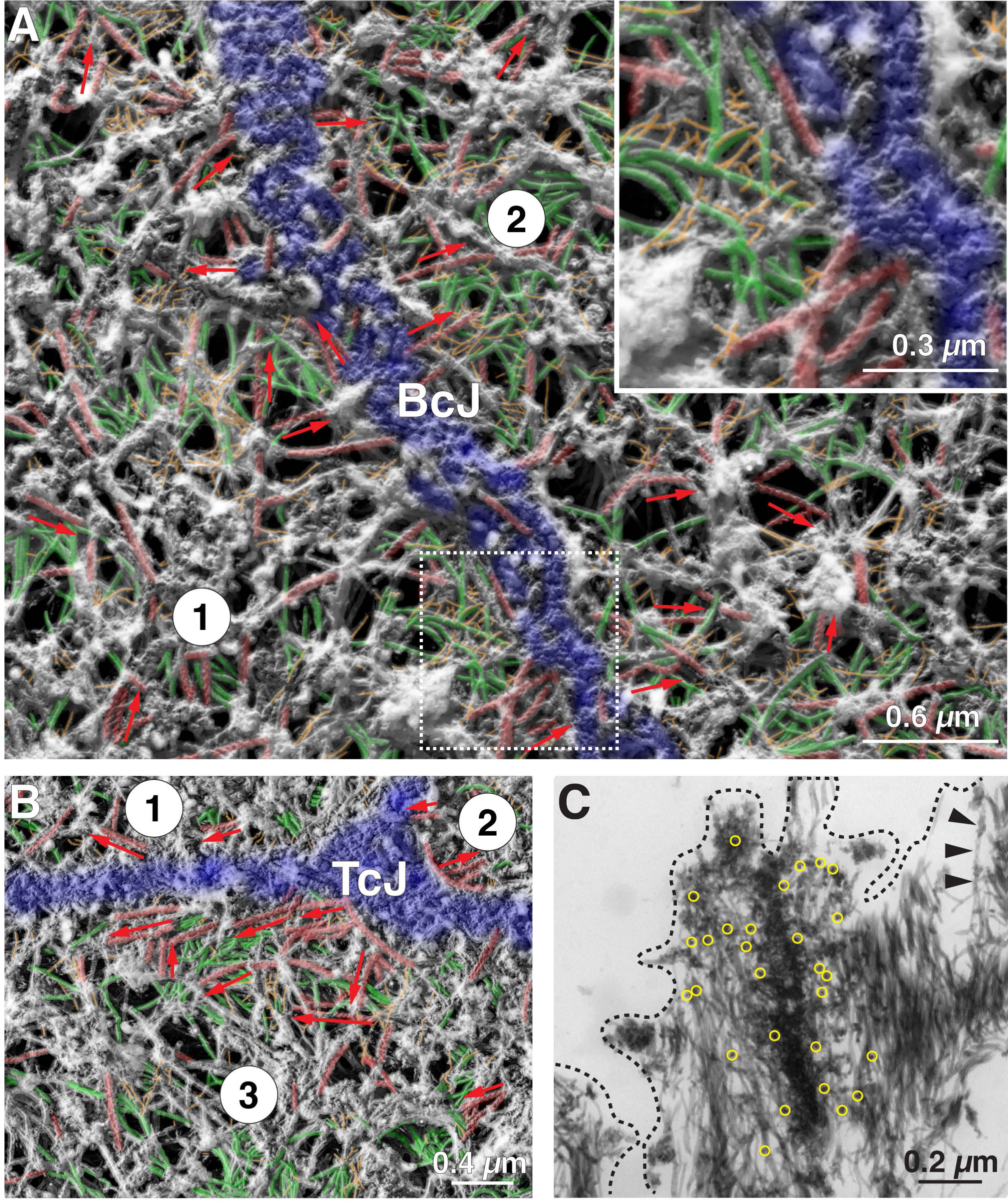
PREM analysis of umbrella cell cytoskeleton near the AJC. **(A)** Cytoskeleton near the bicellular junction (BcJ) was assessed using PREM. The junction itself is false-colored blue, actin is colored red, keratin filaments are green, and cytolinkers are colored orange. Unidentifiable filaments lack false coloring. Red arrows indicate the orientation of actin filaments with the tip of the arrow indicating the pointed end of the S1-decorated filament. The boxed region is magnified in the inset. **(B)** Cytoskeleton network near the tricellular junction (TcJ). **(C)** Thin section TEM of cytoskeletal network near the AJC. Localization of TJP1 (identified using 12nm gold-conjugated secondary antibodies) is indicated by yellow circles. Black arrowheads indicate an S1-decorated actin filament. The approximate position of the plasma membrane is indicated by the black dashed line.

Given the AJC-associated actin ring and the peripheral band of KRT20 identified by confocal microscopy, it was surprising that in these PREM studies there was no obvious concentration of actin or keratins near the AJC. However, PREM only allows for a surface view of the sample. To explore the possibility that the AJC-associated peripheral band of keratins and belt of actin was deep to the surface, we examined cross-sections of PREM processed samples (but lacking the final platinum coating) using TEM. This revealed large numbers of 11-nm keratin filaments sometimes extending up to 500nm on either side of the detergent-resistant, electron-dense core of the tight junction (Fig. 8C). The latter was identified using antibodies to TJP1. The nature of this electron dense core was not explored further. Although, these cross-sectioned samples were also S1 treated, we did not observe decorated actin filaments associated with the tight junction either because they were not resolvable from the dense accumulation of intermediate filaments, or because they were oriented in a manner that did not allow their identification. However, decorated actin filaments (identified by their sympodial branching appearance in thin-section EM) could be found in proximity to the AJC (see black arrowheads in Fig. 8C). These studies revealed that the AJC-proximal band of keratin was closely allied with the AJC including the tight junction; however, the nature of AJC-associated actin was not resolved using these tools.

We also used PREM to explore those regions of apical cytoskeleton away from the AJC. In samples with optimal detergent extraction (i.e., with minimal residual protein contamination), we observed that the bordering elements defining each tessera were composed of filamentous actin, often arranged in parallel bundles of 2 or more actin filaments (Fig. 9A and short arrows in Fig. 9B). While it appeared that the actin filaments were floating above the mesh of subjacent cytoskeletal elements, cross-sections through the extracted network revealed that the actin filaments were intimately associated with underlying keratins (Fig. 9C). A small mass of electron dense material of unknown composition was observed at these points of interaction (yellow asterisks in Fig. 9C). We believe that the keratin filaments below the actin bordering elements revealed by TEM correspond to the larger mesh-forming keratin filaments we identified in our immunofluorescence analysis as they have a similar organization, and they encase areas of similar magnitude (Fig. 4C-D). Unfortunately, we were unable to localize the actin-rich bordering elements using phalloidin, as it does not appear to reliably label these structures. However, we did observe very close interactions between the well-known actin crosslinking protein ACTN4 (α-actinin IV) and KRT20 (Supplementary Fig. S3). Interestingly, the ACTN4 staining was associated with the AJC-associated actin belt, but KRT20 was not. Furthermore, ACTN4^+^ punctae were often spaced along the length of associated KRT filaments, possibly indicating a staggered association of actin with the KRT cytoskeleton.

**Fig. 9.**
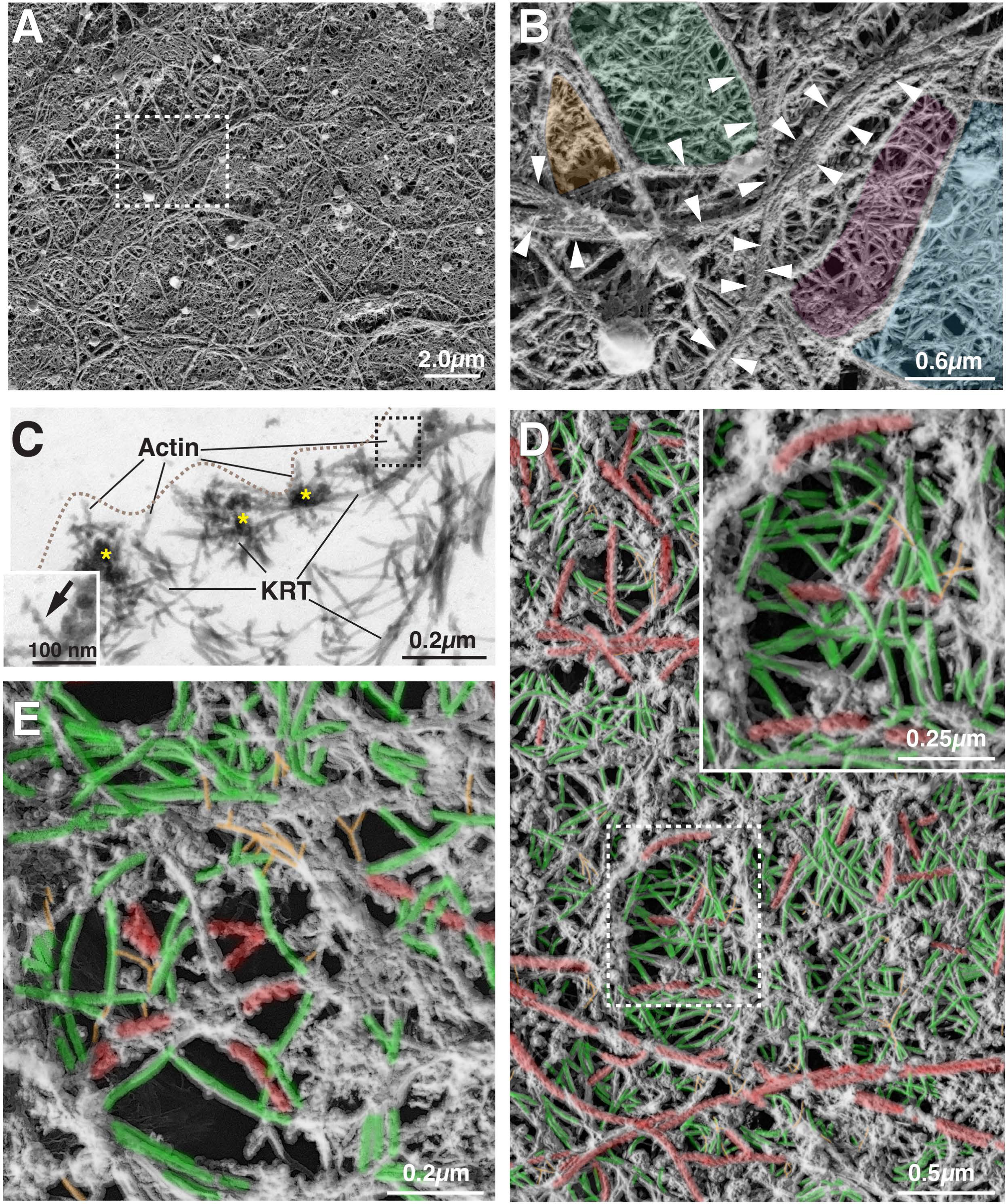
Organization of umbrella cell apical cytoskeleton network revealed by PREM. **(A-B)** Tile-like organization of keratin network. The actin filaments that bordered tesserae were often arranged in parallel bundles of 2 or more actin filaments (indicated by short white arrows in panel B). Example tesserae are false colored in panel B. **(C)** TEM cross section through tesserae revealing the associated actin filaments (facing the apical membrane, which is marked by the dashed line) and subjacent keratin (KRT) filaments. Electron dense material connecting these two filament systems are indicated by yellow asterisks. Boxed region shows an example of an S1-decorated actin filament. **(D-E)** Cytoskeletal mesh lacking lacunae (D) or containing lacunae (E) revealed by PREM. Actin is colored red, keratin filaments are green, and cytolinkers are colored orange. Unidentifiable filaments lack false coloring.

The keratin filaments in the central portions of the tessera were observed to form a crisscrossed or woven pattern extending for short runs and then apparently terminating near actin filaments or other elements not readily discriminated by our analysis (colored green in Fig. 9D). Individual actin filaments were observed interspersed throughout the adjacent cytoskeletal elements (colored red in Fig. 9D), as were cytolinkers that connected the keratin filaments to one another and adjacent bordering elements (colored orange in Fig. 9D). The organization of cytoskeletal elements in lacunae is shown in Fig. 9E. The walls of the lacunae were formed of keratin filaments (sometimes curved), along with short actin filaments, cytolinkers, and unidentified bordering elements. Individual actin filaments and keratins along with cytolinkers were observed to intersect the opening of lacunae. This observation is consistent with a previous report that small numbers of actin and intermediate filaments are closely allied with DFVs in freeze deep-etch preparations ^50^.

In summary, our ultrastructural analysis reveals that the apical cytoskeleton of umbrella cells is organized in a tile-like pattern, with tesserae bordered on their edges by actin filaments and subjacent keratin filaments and filled with crisscrossed/woven keratin filaments crosslinked by cytolinkers. Tesserae are interrupted by lacunae, which likely house unfused DFVs. Finally, the peripheral band of keratin identified in confocal studies was revealed to be a mass of keratins that accumulated on either side of the tight junction, but also extended deep to the desmosomes.

### Impairment of PLEC function causes keratin and actin reorganization, and loss of cell cohesion

Given the importance of cytolinkers such as PLEC in organizing keratin networks ^38, 48^, coupled with the presence of PLEC-like structures in our PREM studies, we sought to determine if PLEC was involved in organizing the native umbrella cell keratin network. Using confocal-SRIP, coupled with 3D surface reconstruction, we identified two general pools of PLEC in umbrella cells. The first one was positioned above the DSG2-labeled desmosomes where it concentrated along either side of the AJC-associated actin ring (colored green in Fig. 10A; Video 6). Furthermore, we observed at sites of cell-cell adhesion, the AJC-associated band of KRT20 was positioned to either side of the AJC-associated PLEC, forming a pentalaminar KRT20-PLEC-actin-PLEC-KRT20 structure (Fig. 10B). The second pool of PLEC was distributed across the apical pole of the cell, where it appeared to be closely associated with the KRT20 mesh (non-AJC PLEC is colored dark blue in Fig. 10A-B). This is consistent with our PREM studies in which we observed PLEC-like cytolinkers that were associated with the mesh layer of keratins. Unfortunately, we were unable to immunolocalize PLEC in PREM samples, but as noted above the cytolinkers we observed by PREM bore the hallmarks of PLEC ^48^.

**Fig. 10.**
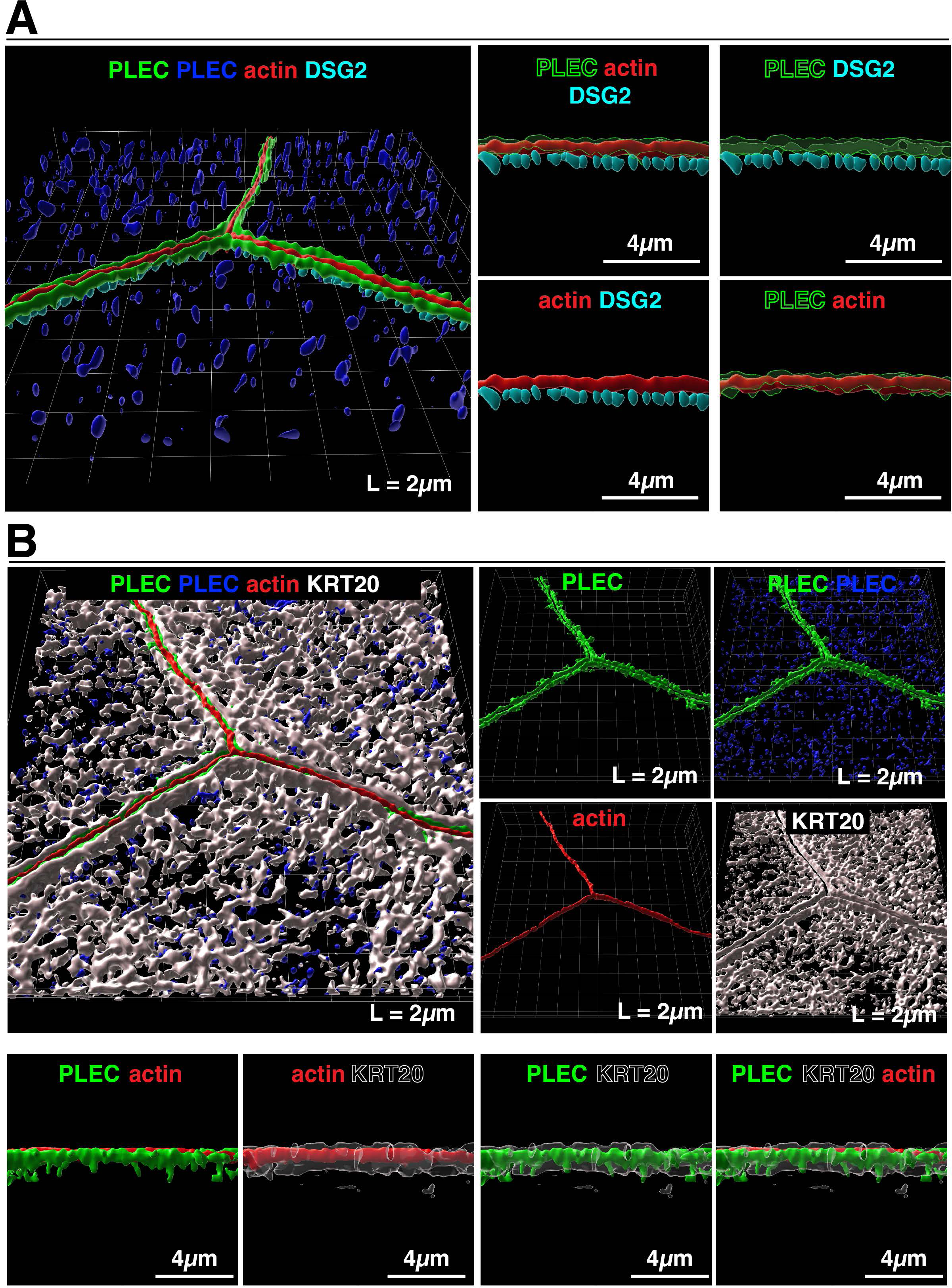
Distribution of PLEC in umbrella cells from filled bladders. **(A)** Left-most image: 3D surface reconstruction of PLEC, AJC-associated actin, and DSG2-labeled desmosomes using data from a confocal-SRIP Z-stack. PLEC signal associated with the AJC is colored green and that associated with the KRT20-labeled mesh is colored dark blue. Right most images: The larger image to the left was rotated along its lateral axis and digitally sliced allowing a head-on view of a bicellular region of the AJC. **(B)** Upper left image: relationship of PLEC to the KRT20 mesh. Upper-right images: signal for the indicated markers are displayed. Lower images: the larger image in the upper left was rotated along its lateral axis and digitally sliced allowing a head-on view of a bicellular region of the AJC. **Video:** A companion video shows the 3D organization of the plectin cytolinker with respect to the AJC (Video6).

PLEC binds keratins by way of a C-terminal intermediate filament-binding domain but can also bind to actin via an N-terminal actin binding domain (see Fig. 11B)^51, 52^. Thus, we hypothesized that PLEC may serve to organize the AJC-associated band of keratin by linking it to the actin ring. As an initial test of this possibility, we asked whether the actin disrupting agents cytochalasin D (CytoD) or the actin-depolymerizing agent latrunculin A (LatA) impacted the organization of the keratin and PLEC networks in stretched bladder samples. CytoD treatment resulted in the accumulation of large aggregates of actin throughout the umbrella cell cytoplasm (Supplementary Fig. S4); however, the AJC-associated actin ring remained intact. PLEC was recruited to a subset of the CytoD-induced actin aggregates (see white arrows in Supplementary Fig. S4), and PLEC staining at the AJC appeared somewhat broader and less tight than that observed in control, DMSO-treated cells (compare PLEC vs DMSO in Supplementary Fig. S4). Despite these effects, CytoD treatment had no obvious impact on the peripheral band of keratin or the overall apical mesh. Surprisingly, LatA (used at concentrations of 1-50 µM) neither affected the actin cytoskeleton nor did it alter the localization of PLEC or KRT20. PLEC also binds to and organizes the microtubule cytoskeleton, in some cases recruiting microtubules to the AJC. In umbrella cells, microtubules were found embedded in the keratin mesh (Supplementary Fig. S5A), and non-AJC PLEC appeared to be in close apposition to microtubules (Supplementary Fig. S5B), but few microtubules were associated with the AJC (Supplementary Fig. S5C). Treatment with the microtubule depolymerizing-agent nocodazole had no visible effect on PLEC or the keratin cytoskeleton (Supplementary Fig. S4).

**Fig. 11.**
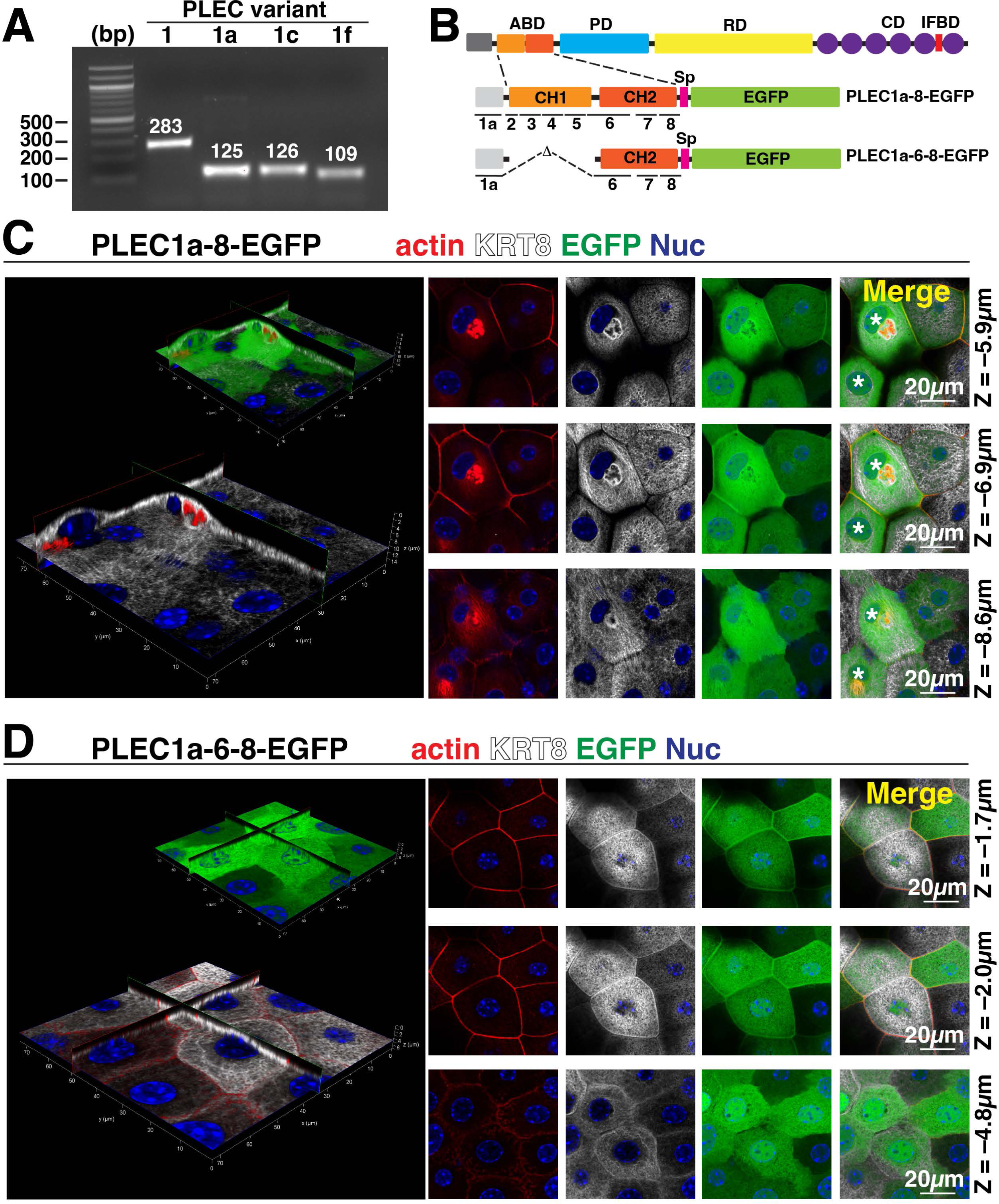
Impact of expressing a dominant negative PLEC construct on the umbrella cell keratin cytoskeleton. **(A)** Use of PCR to assess expression of the indicated PLEC mRNA splice variant in mouse urothelium. The expected size of the amplicon (in bp) is indicated in white text. **(B)** Domain structures of PLEC and the PLEC constructs used in these experiments. Numbers below the constructs indicate the approximate location of the exon boundaries of the *Plec1A* variant. Legend: ABD, actin-binding domain; CD, C-terminal plectin-repeat domain; CH1/2, calponin homology domain 1/2; EGFP, enhanced green fluorescent protein; IFBD, intermediate filament binding domain; PD, plakin domain; RD, coiled-coil rod domain; Sp, spacer (flexible linker). **(C-D)** Mouse bladders were transduced with adenoviruses encoding a dominant-negative PLEC1a-8-EGFP construct (C) or the control PLEC1a-6-8 construct (D). The impact of expression on the actin cytoskeleton and KRT8 cytoskeleton was assessed using confocal microscopy. Larger, left-most images: 3D reconstruction of a confocal Z-stack dissected using orthogonal slicers. Expression of the EGFP-tagged construct is shown in the smaller image in the upper right of the panel. Smaller images: Individual X-Y sections taken at the indicated distance along the Z-axis (top-most image set as Z = 0). The white asterisks in the merged panel for C are examples of two cells that expressed high levels of the construct and exhibit a juxtanuclear aggregate of actin.

We also used a dominant negative approach to assess PLEC-actin interactions. PLEC has multiple N-terminal splice variants, which regulate the recruitment of the cytoskeleton to distinct cellular compartments (e.g., nucleus, focal adhesions, hemidesmosomes)^38, 53, 54^. PLEC1, PLEC1a, PLEC1c, and PLEC1f variants are reportedly expressed by epithelial cells ^53^, and in MDCK cells PLEC1a and PLEC1f were recently shown to be integral to recruitment of rim-associated keratins and the organization of desmosomes ^35^. We used RT-PCR to confirm expression of *Plec1*, *Plec1a*, *Plec1c*, and *Plec1f* variants in mouse urothelium and rat umbrella cells (Fig. 11A and Supplemental Fig. S6). Using this information, we generated two constructs. The first was PLEC1a-8-EGFP, a construct that encoded exon1a of *Plec* along with its complete ABD (encoded by exons 2-8) fused to EGFP (Fig. 11B). This construct was expected to act in a dominant-negative fashion by competing for binding of endogenous PLEC to the actin cytoskeleton. The second construct was PLEC1a-6-8-EGFP, a nominal control construct, which lacked the first calponin homology domain and is reportedly inefficient at binding actin (Fig. 11B)^53,55^.

Because of ease of transduction, we used mouse bladders in the following studies. Note that transduced cells maintain expression of differentiation markers such as uroplakins and achieve a high transepithelial resistance; however, they only have a single nucleus ^7, 56^. The latter is a function of detergent treatment and not a response to viral transduction *per se*. Upon transducing the umbrella cell layer with the PLEC1a-8-EGFP-encoding virus, we observed the expected recruitment of the expressed construct to the region of the AJC (Fig. 11C), but we also observed a filamentous pool of the construct that was just apparent above the background signal (most apparent in the bottom panel, EGFP signal, Fig. 11C). The actin cytoskeleton in these cells was perturbed including the appearance of actin filaments in the cytoplasm near the base of the cells. Intriguingly, the AJC-associated keratin band was absent in cells expressing this construct (Fig. 11C), although the apical keratin mesh had a generally normal appearance. In addition, we noted the appearance of keratins at the points of cell-cell adhesion that were straight and in alignment with filaments on the adjoining cell (Z = –8.6µm in Fig. 11C), consistent with the cells being under a mechanical load. A small fraction of cells (∼ 20%) was observed to express large amounts of this construct and these cells formed a juxtanuclear cytoplasmic aggregate that was actin and KRT20 positive (cells marked with an asterisk in right-most panels of Fig. 11C). We also observed a loss of PLEC staining near the AJC in cells expressing this construct, but not in adjacent un-transduced cells (Supplementary Fig. S7). The control PLEC1a-6-8-EGFP construct had an overall cytoplasmic appearance, although some of it was recruited to the region of the AJC (Fig. 11D). If this reflects binding via sequences encoded by PLEC exon 1a, or weak adhesion to the actin ring was not assessed. However, in cells expressing this construct we did not observe the development of actin-rich cytoplasmic aggregates or actin fibers, and the apical keratin mesh and band of keratin near the AJC were not affected in these cells (Fig. 11D). Furthermore, we noted minimal loss of PLEC staining at the AJC of cells expressing this control construct (Supplementary Fig. S7).

To further explore a functional role for PLEC in the organization of the umbrella cell keratin cytoskeleton we turned to plecstatin-1, a newly described anti-tumor drug that selectively targets PLEC and which mimics the effects of *Plec* knockout in MDCK cells ^35, 40^. When the urothelium was treated with 25µM plecstatin-1, we noted cells with focal abnormalities in the organization of their keratin and actin cytoskeletons. These changes included the formation of actin- and PLEC-rich foci that formed in the central portions of the cells or at sites along the bi-cellular and tri-cellular junctions (Fig. 12). These foci exhibited a corresponding decrease in KRT20 signal, leaving discontinuities in the keratin network including loss of the AJC-associated keratin band in affected cells (see white asterisks in merged panel of Fig. 12). These lesions were also accompanied by discontinuities in the tight junction (marked by CLDN8; Supplementary Fig. S8A) and loss of cell-cell cohesion (yellow arrows in Fig. 12). In some case, we observed regions where umbrella cells were missing, revealing the underlying intermediate cells (see dashed region in the upper row, left-most panel of Supplementary Fig. S8A). When plecstatin-1-treated cells were examined by PREM, the foci were revealed to be mesa-like structures that were often elevated and had a flat top (Supplementary Fig. S8B). Surrounding the mesa were accumulations of woven actin, as well some keratins and cytolinkers (Supplementary Fig. S8C). Taken together, these studies reveal that PLEC function is integral to the organization of the umbrella cell keratin cytoskeleton, and that disruption of PLEC function leads to formation of mesa-like foci that are characterized by focal disassembly of the keratin mesh, loss of the AJC-proximal band of keratin, compensatory changes in the actin cytoskeleton, AJC dysfunction, and loss of umbrella cell cohesion.

**Fig. 12.**
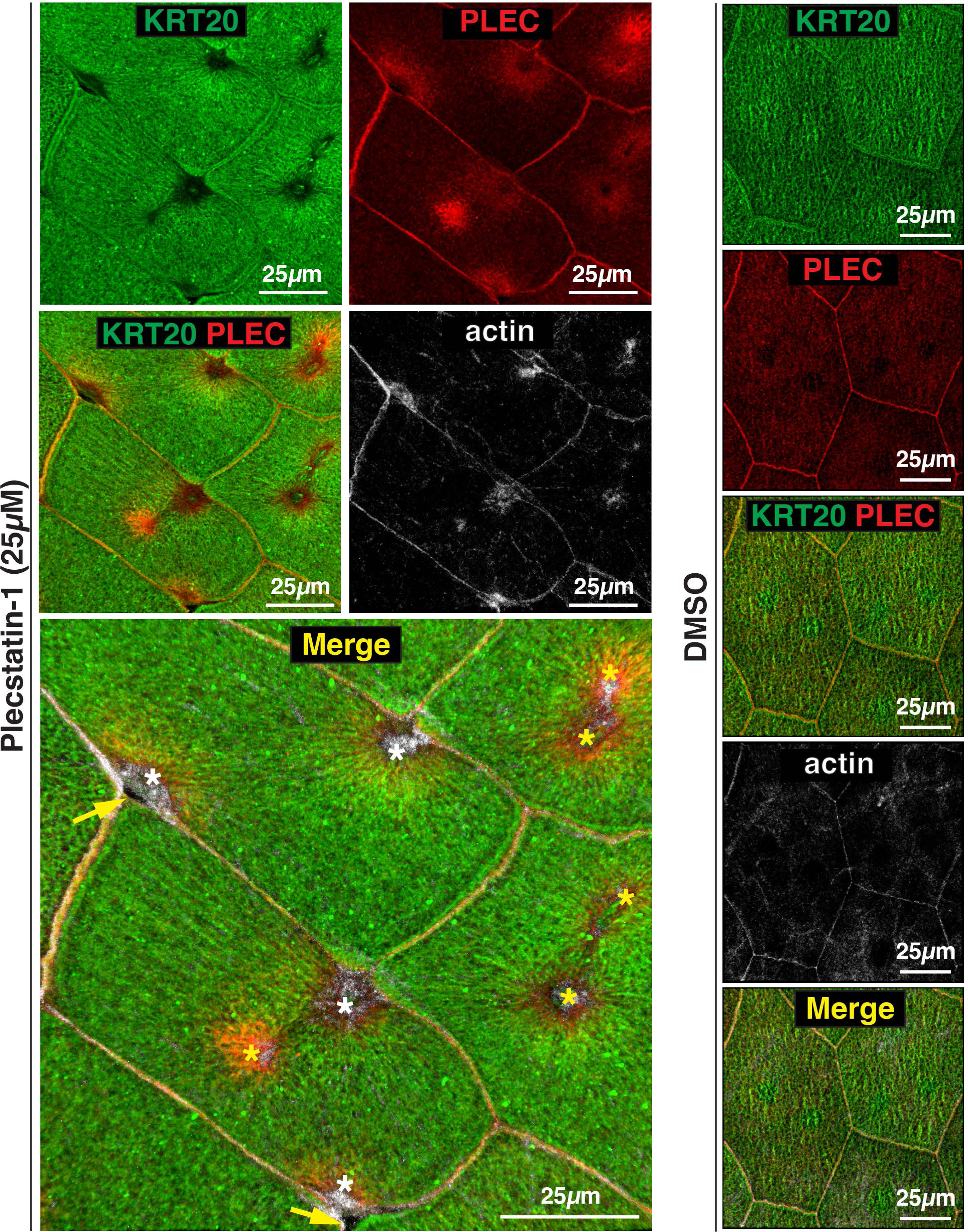
Effect of plecstatin-1 on the cytoskeletal network of umbrella cells. Stretched bladder tissue was treated with 25 µM plecstatin-1 or DMSO solvent for 60 min at 37°C. A 3D reconstruction of a Z-volume obtained using confocal microscopy was generated. In the merged panel, yellow arrows indicate regions of lost cohesion, white asterisks are examples of mesa-like foci forming near the AJC, and yellow asterisks are mesa-like foci forming in the central regions of the cell.

## DISCUSSION

Key contributors to epithelial stability and cohesion are desmosomes and the associated keratin cytoskeleton ^10, 13^. Yet, we have few insights into the organization of the umbrella cell keratin cytoskeleton, the adaptations that the umbrella cell keratin cytoskeleton makes in response to bladder filling, or the molecules that regulate this network. Our studies reveal the following: (i) that umbrella cells have an elaborate apical keratin cytoskeleton organized as a tile-like mesh; (ii) that bladder filling and attendant increase in wall tension leads to a reorganization of the umbrella cell desmosome/keratin network including formation of a girded layer; (iii) that the plectin cytolinker is integral to the continuity of the umbrella cell keratin cytoskeleton and PLEC dysfunction leads to focal keratin loss and umbrella cell de-adhesion. Taken together, these observations support the likelihood that the umbrella cell keratin cytoskeleton contributes to umbrella cell mechanical stability and function during the bladder filling/voiding cycle.

The keratin networks of epithelial cells such as HaCaT keratinocytes, MDCK cells, native retinal pigment epithelial cells, and blastocysts are organized as loose “fishnet-like” meshes ^30, 32^. Integral to these meshes, is a rim-and-spoke architecture in which thicker keratin filaments (i.e., spokes) connect desmosomes to the nucleus and fine keratin filaments (i.e., rims) interlock the desmosomes ^30^. Other patterns of keratin organization exist. For example, in enterocytes, keratins are observed to accumulate at the apical pole of the cell where they are observed to crosslink the actin-rich rootlets of microvilli ^57, 58^, and during mouse airway development, keratins form a thin mesh that surrounds the basal bodies of ciliated airway cells ^31^. In contrast, the apical keratin cytoskeleton of native umbrella cells is organized as a dense tile-like mesh. This mesh is comprised of bordering elements filled with short, crisscrossing and/or woven keratin filaments, which are further crosslinked by cytolinkers including PLEC. The bordering elements are comprised, in part, of cortical actin filaments that are not readily detected using phalloidin but are visible by PREM after S1-decoration. The lack of phalloidin staining may reflect the thin nature of these actin filaments (typically 2-3 actin filaments in a side-by-side configuration), which may not be easily detected or resolved by confocal-SRIP microscopy. Alternatively, their conformation or associated proteins may prevent phalloidin binding as has been described in other contexts ^59–61^. Below these actin filaments are subjacent keratin filaments, which based on their organization and dimensions likely give the keratin network its mesh-like appearance when viewed *en face*. Punctuating this mesh are lacunae, which house DFVs prior to fusion ^36^.

Given the need for the umbrella cell layer to withstand pressures climbing as high as ∼ 60-80cm H2O during voiding contractions ^62^, we anticipate the need for a semi-flexible keratin network that can stand up to distension, that allows for exocytosis, and that is able to return to its ground state within minutes of voiding. Several characteristics of the umbrella cell network we describe fits these needs. A critical contributor to the mechanical properties of fibrous networks is the mechanical behavior of their filaments including their elasticity and their stiffness, which is described by the relationship between their contour length (Lc) and persistence length (Lp; i.e., bending stiffness) ^63, 64^. In general, a stiff fiber is characterized by LP >> LC, whereas fibers that are semi-flexible have an LP ≅ LC, and flexible fibers have an LP << LC. Given the reported persistence length of keratins (∼ 0.3 µm) ^65^, and the contour length of keratins within tesserae (∼ 0.5 µm), these fibers can be classified as semiflexible. However, the presence of crosslinks (e.g., those mediated by PLEC) would alter the LP/LC relationship because LC is measured between crosslinked segments and when LP > LC, a less flexible fiber is formed. Thus, by altering PLEC crosslinks, one could regulate the stiffness of the umbrella cell keratin network. Longer keratin filaments, such as those subjacent to actin bordering elements, may be more flexible (as LP << LC). In contrast, the cortical actin bordering elements may serve to stiffen the overall network as the Lp of actin is ∼ 18 µm ^66^, and the length of the actin bordering elements is variable but on the order of 1-2 µm (thus LP >> LC). However, the association of these cortical actin filaments with the umbrella cell apical membrane may result in softening as is described for spectrin networks in erythrocytes ^67^. Likewise, the short actin filaments (typically <1.0 µm) we observed interspersed throughout the network, in lacunae, and near the junctions may also serve to stiffen the network. Other critical features of fibrous networks are parameters associated with network structures including filament density, which regulates the rigidity of the network, crosslinks, which govern the network stability, and orientation of filaments, which affects mechanical anisotropy (material characteristics that depend on the direction of mechanical load) ^64^. Further study will be needed to understand the mechanics of the umbrella cell keratin network and how it contributes to umbrella cell physiology.

The other prominent component of the apical keratin network we describe is a thin band of keratin, originally described as a “frame” ^36^, that abuts the AJC and which we revealed by confocal-SRIP microscopy and thin-section EM to be an accumulation of keratin filaments on either side of the tight junction and AJC-associated actin ring. The function(s) of this keratin band is unknown, but it could serve as a reserve of keratin filaments that would allow the umbrella cell to expand during bladder filling (and then function as a site of storage after voiding), or it may be a site of keratin nucleation/assembly. The keratin band does not obviously function as a “rim” in the sense that it is positioned above the desmosomes and instead appears to be organized by the PLEC cytolinker, which is sandwiched between the AJC-associated actin ring and the keratin band. Thus, in addition to previous evidence that PLEC regulates rim keratins, spokes, and associated desmosomes in MDCK cells ^35^, our data reveals that in native umbrella cells PLEC organizes the overall keratin network as well as the AJC-associated keratin band and its association with the actin cytoskeleton.

We also observed a possible role for PLEC in regulating AJC continuity (the actin ring and tight junction in particular). In plecstatin-1-treated umbrella cells, we observed focal loss of the AJC-associated actin cytoskeleton and a corresponding loss of tight junction continuity, eventually leading to loss of cohesion. Given the importance of actin in tight junction and adherens junction function ^68^, the loss of actin likely contributes to the eventual disruption of cell cohesion and cell detachment we observed. Our data comports with evidence that conditional deletion of *Plec* in intestinal epithelial cells (enterocytes) leads to the formation of aberrant tight junctions, adherens junctions, and desmosomes, leading to intercellular gaps and a leaky gut phenotype ^42^. Furthermore, disruption of tight junctions in biliary tree cholangiocytes is observed in conditional hepatocyte/cholangiocyte *Plec* deleted mice ^41^.

Our studies also revealed several adaptations that the umbrella cell’s keratin network makes during bladder filling and which we believe ensures cell continuity. The first adaptation is related to the reported thinning of the bordering elements of the keratin network mesh observed in filled bladders ^36^. This may indicate that the keratin mesh can stretch during bladder filling; however, given the large increase in cell perimeter during filling, along with the introduction of new desmosomes to the expanding AJC, it is likely new keratin mesh is synthesized or assembled during the bladder filling phase. The second adaptation is expansion of the desmosome necklace to further accommodate the increase in surface area and change in cell shape, as well maintain tissue cohesion. Given our previous studies of the tight junction and adherens junctions ^7^, it is likely that this expansion of the desmosomal necklace is driven in part by actin polymerization and exocytosis of junction components such as DSG2 and DSC2. However, it is possible that it may also involve the reversible recruitment of desmosomes from the basolateral surfaces of the umbrella cell. In contrast, shrinking of the cell perimeter in response to voiding would likely entail non-muscle myosin II-mediated contraction and endocytosis of desmosomes, which has been studied in other settings ^69–72^.

The third adaptation is the ability of the keratin network to support the exocytosis of large numbers of subapical DFVs in response to filling. An emerging literature indicates that intermediate filaments can regulate vesicular traffic ^73^. We previously reported that DFV maturation likely involves a Rab11a-Rab8a cascade whereby Rab11a^+^ DFVs, which predominate in the keratin network, mature into a release-ready pool of Rab8^+^ DFVs (and a separate subpool of Rab27b^+^ DFVs) that accumulate below the apical membrane ^74, 75^. Given previous reports of actin and keratin association with DFVs in lacunae ^36, 50^, it is possible that these cytoskeletal elements promote DFV maturation. Moreover, the elastic nature of keratin filaments would also allow rapid and facile penetration of the keratin mesh in the situation whereby DFVs undergo massive exocytosis ^5^.

The fourth adaptation, and perhaps the most striking change we observed in response to filling, is the formation of a girded layer as umbrella cells transition from a “cuboidal” morphology to a squamous one. Comprised of nested ovals of keratin filaments, the girded layer is a site of keratin attachment to the desmosomes at the AJC, but also those desmosomes at the umbrella cell:intermediate cell junction. The latter would ensure overall urothelial cohesion by strongly adhering umbrella cells to subjacent intermediate cells. We further observe that the keratin ovals are held together by struts formed from straight-appearing keratin filaments that cross the ovals at ∼90° angles. This organization is consistent with a cell reacting to forces generated along its entire perimeter as the bladder fills. We further observed that the apical pole umbrella cell nuclei are integrated into this girded layer, with close apposition of keratin filaments with the nuclear ectocytoskeleton. This may indicate a mechanism to convey cell-associated changes in cell shape and mechanical stress to the nucleus. The girded layer becomes more prominent when the bladder is filled to capacity, additional evidence that this structure likely plays an important role in maintaining umbrella cell cohesion during filling. Furthermore, the girded layer is absent in the umbrella cells of quiescent bladders. Thus, umbrella cells can rapidly adapt to a changing mechanical environment by altering the organization of its keratin network in response to filling, and then rapidly reverse these changes upon voiding.

### Summary

Epithelial cells must maintain a cohesive barrier in the face of mechanical forces such as when the skin is stretched, the lungs fill with air, food passes through the gut, or as the bladder fills with urine. In the case of umbrella cells, previous studies revealed mechanisms that contribute to tissue continuity during bladder filling including cell shape changes, surface area changes, and expansion of the tight and adherens junction ^1, 2, 7^. Our current studies reveal that the umbrella cell keratin network, organized as a tile-like mesh, has several features that would contribute to umbrella cell mechanical stability including stretching and/or growing their keratin network, expanding their AJC-associated desmosomal necklace, and by forming a girded layer in response to filling, changes which are rapidly reversed upon voiding.

## MATERIALS and METHODS

### Animals

Female Sprague-Dawley rats, 10-11wks old (225-250g in weight), were obtained from Inotiv-Envigo (West Lafayette, IN). They were housed, two animals per cage, in solid-bottom plastic cages. They were maintained under a 12-h day/night cycle and were fed standard rat chow and given water *ad libitum*. Female C57Bl6/J mice, 8-10wks in age, were obtained from the Jackson Laboratory (Bar Harbor, ME). Mice were group housed (up to five mice/cage) in solid-bottom plastic cages under a 12-h day/night cycle. They were fed standard mouse chow and given water *ad libitum.* Animal studies were performed in accordance with relevant guidelines/regulations of the Public Health Service Policy on Humane Care and Use of Laboratory Animals and the Animal Welfare Act, and under the approval of the University of Pittsburgh Institutional Animal Care and Use Committee. Rats and mice were euthanized by CO2 inhalation, followed by thoracotomy as a secondary method.

### Reagents and Antibodies

Unless specified otherwise, all chemicals were obtained from Sigma-Aldrich (St Louis, MO). The source of primary and secondary antibodies used in this study are detailed in **Table 1**. Rhodamine-labeled phalloidin and DAPI were obtained from Molecular Probes (ThermoFisher Scientific, Grand Island, NY) and Phalloidin-405 from Abcam (Waltham, MA).

**Table 1:**
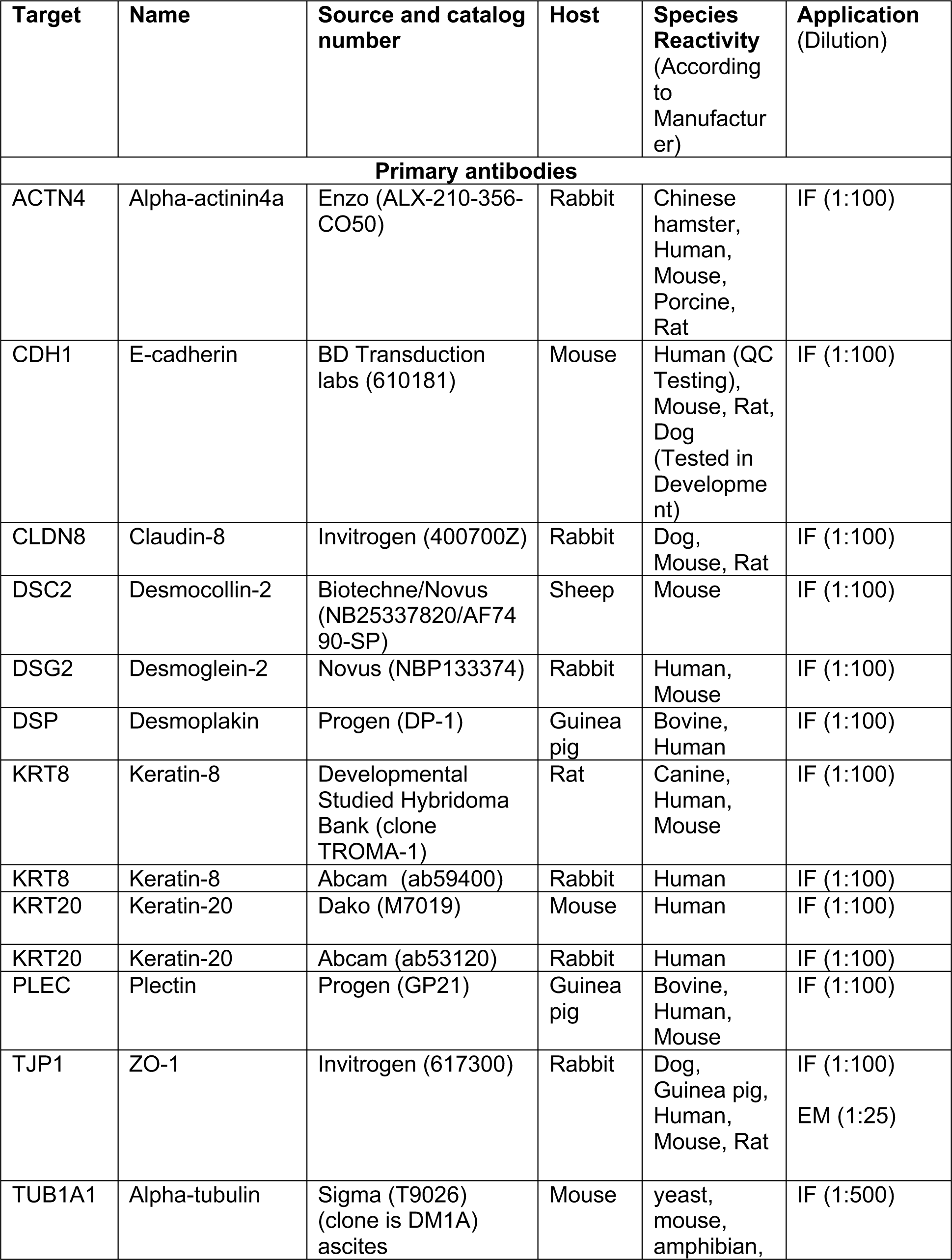
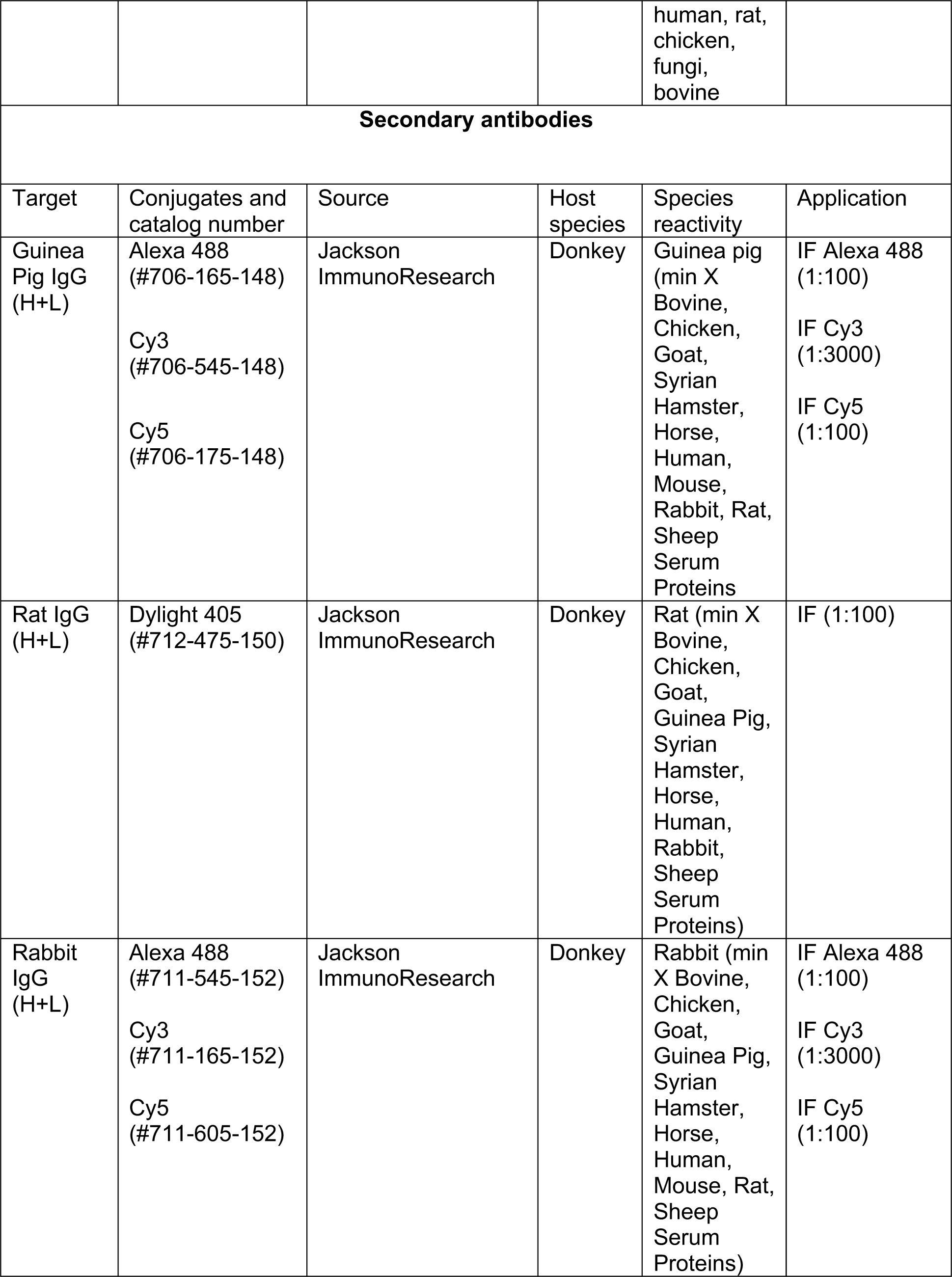
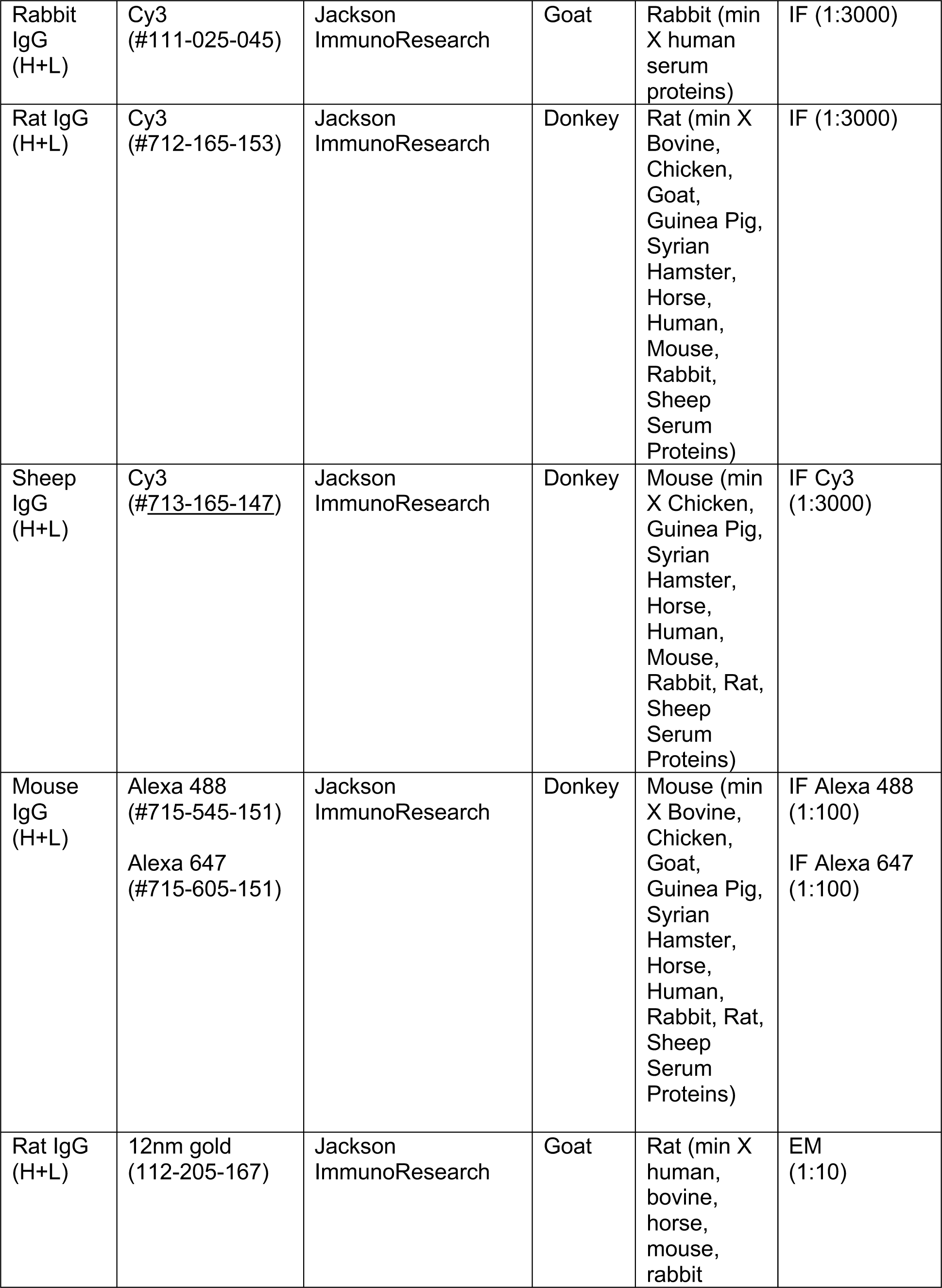
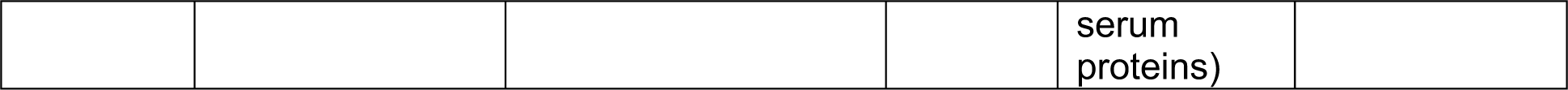
Antibodies used in this study.

| Target | Name | Source and catalog number | Host | Species Reactivity (According to Manufacturer) | Application (Dilution) |
| --- | --- | --- | --- | --- | --- |
| **Primary antibodies** |  |  |  |  |  |
| ACTN4 | Alpha-actinin4a | Enzo (ALX-210-356-CO50) | Rabbit | Chinese hamster, Human, Mouse, Porcine, Rat | IF (1:100) |
| CDH1 | E-cadherin | BD Transduction labs (610181) | Mouse | Human (QC Testing), Mouse, Rat, Dog (Tested in Development) | IF (1:100) |
| CLDN8 | Claudin-8 | Invitrogen (400700Z) | Rabbit | Dog, Mouse, Rat | IF (1:100) |
| DSC2 | Desmocollin-2 | Biotechne/Novus (NB25337820/AF7490-SP) | Sheep | Mouse | IF (1:100) |
| DSG2 | Desmoglein-2 | Novus (NBP133374) | Rabbit | Human, Mouse | IF (1:100) |
| DSP | Desmoplakin | Progen (DP-1) | Guinea pig | Bovine, Human | IF (1:100) |
| KRT8 | Keratin-8 | Developmental Studied Hybridoma Bank (clone TROMA-1) | Rat | Canine, Human, Mouse | IF (1:100) |
| KRT8 | Keratin-8 | Abcam (ab59400) | Rabbit | Human | IF (1:100) |
| KRT20 | Keratin-20 | Dako (M7019) | Mouse | Human | IF (1:100) |
| KRT20 | Keratin-20 | Abcam (ab53120) | Rabbit | Human | IF (1:100) |
| PLEC | Plectin | Progen (GP21) | Guinea pig | Bovine, Human, Mouse | IF (1:100) |
| TJP1 | ZO-1 | Invitrogen (617300) | Rabbit | Dog, Guinea pig, Human, Mouse, Rat | IF (1:100)<br>EM (1:25) |
| TUB1A1 | Alpha-tubulin | Sigma (T9026) (clone is DM1A) ascites | Mouse | yeast, mouse, amphibian, | IF (1:500) |

|  |  |  |  | human, rat,<br>chicken,<br>fungi,<br>bovine |  |
| --- | --- | --- | --- | --- | --- |
| **Secondary antibodies** |  |  |  |  |  |
| Target | Conjugates and<br>catalog number | Source | Host<br>species | Species<br>reactivity | Application |
| Guinea<br>Pig IgG<br>(H+L) | Alexa 488<br>(#706-165-148)<br>Cy3<br>(#706-545-148)<br>Cy5<br>(#706-175-148) | Jackson<br>ImmunoResearch | Donkey | Guinea pig<br>(min X<br>Bovine,<br>Chicken,<br>Goat,<br>Syrian<br>Hamster,<br>Horse,<br>Human,<br>Mouse,<br>Rabbit, Rat,<br>Sheep<br>Serum<br>Proteins | IF Alexa 488<br>(1:100)<br>IF Cy3<br>(1:3000)<br>IF Cy5<br>(1:100) |
| Rat IgG<br>(H+L) | Dylight 405<br>(#712-475-150) | Jackson<br>ImmunoResearch | Donkey | Rat (min X<br>Bovine,<br>Chicken,<br>Goat,<br>Guinea Pig,<br>Syrian<br>Hamster,<br>Horse,<br>Human,<br>Rabbit,<br>Sheep<br>Serum<br>Proteins) | IF (1:100) |
| Rabbit<br>IgG<br>(H+L) | Alexa 488<br>(#711-545-152)<br>Cy3<br>(#711-165-152)<br>Cy5<br>(#711-605-152) | Jackson<br>ImmunoResearch | Donkey | Rabbit (min<br>X Bovine,<br>Chicken,<br>Goat,<br>Guinea Pig,<br>Syrian<br>Hamster,<br>Horse,<br>Human,<br>Mouse, Rat,<br>Sheep<br>Serum<br>Proteins) | IF Alexa 488<br>(1:100)<br>IF Cy3<br>(1:3000)<br>IF Cy5<br>(1:100) |
| Rabbit IgG (H+L) | Cy3 (#111-025-045) | Jackson ImmunoResearch | Goat | Rabbit (min X human serum proteins) | IF (1:3000) |
| Rat IgG (H+L) | Cy3 (#712-165-153) | Jackson ImmunoResearch | Donkey | Rat (min X Bovine, Chicken, Goat, Guinea Pig, Syrian Hamster, Horse, Human, Mouse, Rabbit, Sheep Serum Proteins) | IF (1:3000) |
| Sheep IgG (H+L) | Cy3 (#713-165-147) | Jackson ImmunoResearch | Donkey | Mouse (min X Chicken, Guinea Pig, Syrian Hamster, Horse, Human, Mouse, Rabbit, Rat, Sheep Serum Proteins) | IF Cy3 (1:3000) |
| Mouse IgG (H+L) | Alexa 488 (#715-545-151)<br>Alexa 647 (#715-605-151) | Jackson ImmunoResearch | Donkey | Mouse (min X Bovine, Chicken, Goat, Guinea Pig, Syrian Hamster, Horse, Human, Rabbit, Rat, Sheep Serum Proteins) | IF Alexa 488 (1:100)<br>IF Alexa 647 (1:100) |
| Rat IgG (H+L) | 12nm gold (112-205-167) | Jackson ImmunoResearch | Goat | Rat (min X human, bovine, horse, mouse, rabbit) | EM (1:10) |
|  |  |  |  | serum<br>proteins) |  |

### Preparation of stretched bladders, filled bladders, voided bladders, and quiescent bladders

Stretched bladders were prepared as follows. Rats were euthanized and an incision was made along the ventral caudal midline, the bladder was exposed, clamped at the neck region (i.e., adjacent to the urethra) using a hemostat, and then excised. The bladder was carefully cut open along its midline from neck to dome using sharp surgical scissors and then rinsed in Krebs buffer (110mM NaCl, 25mM NaHCO3, 5.8mM KCl, 1.2mM MgSO4, 4.8mM KH2PO4, 11mM glucose, 2mM CaCl2, gassed with 5% v/v CO2). The cut-open bladder was submerged in Krebs buffer and then pinned out on a rubber mat forming a 2 x 2 cm square. The tissue was then allowed to recover for 30min in a tissue culture incubator held at 37°C and gassed with 5% v/v CO2/95% v/v air. The tissue was then drug treated, extracted, and/or fixed as described below. SEM confirmed that tissue prepared in this manner was smooth and flat (i.e., lacked any rugae or surface folds), the umbrella cells were polyhedral and not obviously pulled in one direction, and the tissue was morphologically similar to that observed when bladders are filled to 0.5 ml and then perfusion fixed.

Filled, voided, and quiescent bladders were prepared using our previously described techniques ^7^. Rats were anesthetized using 3% v/v isoflurane and then injected in their dorsal neck scruff subcutaneously with 1.4g/kg urethane prepared fresh in dH2O and sterile filtered through a 0.22μm STERIFLIP-GP filter (Millipore Sigma) prior to use. The isoflurane was maintained for 5min and then stopped. The animals were allowed to reach proper anesthetic depth over a period of ∼ 0.5-1.0h, which was confirmed by lack of a response to a toe pinch. The animals were catheterized by inserting a 22-g Jelco IV catheter (Smiths-Medical, Minneapolis, MN) into the urethra and the animals were subjected to Credé’s maneuver to void their bladders prior to the start of the experiment. Subsequently, animals in the filled and voided groups had their catheter ports closed, and their bladders were filled over 30min to a final volume of 500-1000μL using a Harvard Apparatus (Holliston, MA) PHD ultra syringe pump. Animals in the voided group had their catheter ports re-opened and then allowed to void for5 min. The animals with quiescent bladders remained catheterized, with catheter ports left open, for the full 30-min period, maintaining their bladders in an empty state. At the end of the experiment, animals were perfusion fixed using the following methods. A thoracotomy was performed, the caudal vena cava was cut, and 50ml of 100mM Sorensen’s phosphate buffer, pH 7.4 (phosphate buffer) at 37°C was perfused through the left ventricle using an 18-g needle. Subsequently, the perfusate was switched to phosphate buffer containing 4% w/v paraformaldehyde. The bladders were excised, placed in phosphate buffer containing 4% w/v paraformaldehyde, cut open down their midline and pinned out on a rubber mat, with minimal stretching, to expose the apical-most umbrella cell layer. The tissues were stored at 4°C in 1% w/v paraformaldehyde in phosphate buffer until ready for processing.

### Thin-section transmission electron microscopy

Stretched bladders were fixed for 60min at room temperature in 2.0% v/v glutaraldehyde (EMS; Electron Microscopy Sciences, Hatfield, PA), 4% v/v EM-grade paraformaldehyde in phosphate buffer. Alternatively, we used immuno-labeled PREM samples, prepared as described below. The fixed tissues were cut into 1-2mm^2^ tissue blocks, rinsed with 100mM cacodylate, pH 7.4 buffer and then osmicated in reduced 1.0% w/v OsO4 (EMS) containing 1.5% w/v K4Fe(CN)6 and dissolved in 100mM cacodylate buffer, pH 7.4 for 60min at room temperature. The tissue blocks were rinsed three times with water and *en bloc* stained overnight at 4°C with an aqueous solution of 0.5% w/v uranyl acetate (EMS). The tissue blocks were dehydrated in ethanol, incubated in a 1:1 mixture of propylene oxide:LX112 epon substitute (Ladd Research, Essex Junction, VT) overnight, and then three changes of LX112 medium over the next two days, prior to embedding in flat rectangular silicone molds (Ladd Research) filled with LX112 medium. Samples were cured for 48h at 60° C, the blocks were trimmed, and the tissue sectioned at 70nm using a Leica Microsystems Ultracut R ultramicrotome (Wetzlar, Germany) outfitted with a Diatome Ultra 45° diamond knife (EMS-Diatome, Hatfield, PA). Sections, collected on Butvar-coated nickel grids, were contrasted with uranyl acetate and lead citrate (EMS) and viewed and photographed using a JEM1400 FLASH electron microscope (JEOL, Peabody, MA).

### Cryosectioning of bladder tissue

Fixed, filled, and quiescent bladders were placed dome side down in cryomolds (15 x 15 x 5mm; ThermoFisher Scientific) filled with Optimal Cutting Temperature (OCT) solution (Tissue-Tek, Sakura Finetek, Torrance, CA), and flash frozen by placing the cryomold on a pool of liquid nitrogen. The blocks were stored at -80°C in tightly sealed plastic bags. Cryosections were cut using a CM1950 cryostat (Leica Microsystems, Buffalo Grove, IL; 8-12µm sections; -20°C chamber and -18°C knife temperatures), collected on Superfrost Plus glass slides (ThermoFisher Scientific), and held within the cryochamber at -20°C prior to immunolabeling or stored at -80°C.

### Processing and immunolabeling of whole-mounted and cryo-sectioned bladder tissue

Stretched bladder tissue was prepared as described above and then fixed with 4% w/v paraformaldehyde in phosphate buffer for 30min. The tissue was washed three times with phosphate buffered saline (PBS) for 5min and unreacted paraformaldehyde quenched using PBS containing 0.1% v/v Triton X-100, 20mM glycine, pH 8.0, and 75mM NH4Cl2. The tissue was then treated with Permeabilization and Cytoskeletal Stabilization (PCS) buffer, which was optimized for PREM (see below), and which is comprised of 100mM PIPES, pH 6.9 (titrated with KOH), 1mM MgCl2, 1mM EGTA, 5% v/v poly(ethylene glycol) octyl ether detergent (POE; stored under nitrogen), and 0.2% w/v polyethylene glycol (20,000 mwt; ThermoFisher Scientific) for 45 min at room temperature on a slowly-rotating orbital shaker (30 rpm). The tissue was rinsed with PEM buffer (100mM PIPES, pH 6.9, 1mM MgCl2, 1mM EGTA) three times.

Immunofluorescent labeling of whole-mounted and cryosectioned bladder tissue was performed at room temperature, unless otherwise indicated. For cryosectioned tissue, the slide-mounted sections were washed three times with PBS for 5min and unreacted paraformaldehyde quenched using PBS containing 0.1% (v/v) Triton X-100, 20mM glycine, pH 8.0, and 75mM ammonium chloride. The procedure for whole-mounted or cryosectioned tissue was similar. Samples were washed three times with PBS for 5min, and then three times quickly with Block Solution (which contained 0.7% w/v fish-skin gelatin, 0.025% w/v saponin, and 0.02% w/v sodium azide, all dissolved in PBS). The tissue was then incubated for 30min in Block Solution supplemented with 5% v/v donkey serum, and then incubated overnight at 4°C in Block Solution containing a mixture of primary antibodies. Subsequently, the tissue was washed three times quickly with Block Solution, and then three times for 5min with Block Solution prior to incubating the tissue with secondary antibodies, diluted in Block Solution, for 1h. Tissue was then washed three times quickly with Block Solution, three times for 5min with Block Solution, and then three times with PBS. The samples were post-fixed with 4% (w/v) paraformaldehyde dissolved in phosphate buffer for 10min (tissue sections) or 20min (whole-mount), after which the tissue was washed three times with PBS. After labeling, cryosections were covered with a drop of Slowfade Diamond Antifade mountant (refractive index of 1.42; ThermoFisher Scientific), covered with a Gold Seal Cover Glass (number 1.5, ThermoFisher Scientific), and sealed around the edges with a thin layer of clear nail polish (EMS). Whole-mount tissue was placed, mucosal surface facing up, within a square well created with nail polish in which a drop of Slowfade Diamond Antifade Mountant was added. An additional drop of mountant was added to the top of the tissue. A cover glass was placed over the tissue and sealed around its edges with a thin layer of nail polish. Samples were stored at -20°C. Use of Slowfade-Glass antifade mountant (ThermoFisher Scientific), which has a refractive index of 1.52, did not significantly enhance our image quality and was not routinely employed.

### Image acquisition, 3D image reconstruction, and video production

Images were captured by confocal microscopy using a Leica DMI8 microscope and either a Leica HC PL APO CS2 10X, 0.4NA dry objective, a Leica HC PL APO CS2 40X, 1.30NA oil objective, or a Leica HC PL APO CS2 63X, 1.4NA oil objective and the appropriate laser lines of a Leica Microsystems SP8 Stellaris confocal system outfitted with a 405nm laser diode and a white-light laser. The signal from the HyDS and HyDX detectors was optimized using the Q-LUT option and crosstalk between channels was prevented by use of spectral detection coupled with sequential scanning. For standard confocal microscopy, 8-bit images (1024 x 1024) were collected at 600Hz using 3-line averages, with a pinhole diameter of 1.0Airy unit, and system-optimized parameters for the Z-axis. For SRIP-confocal microscopy we used the LIGHTNING deconvolution processing module set to its highest quality setting. Images (16 bit), captured using a pinhole set to 0.5Airy units, were collected using system optimized settings for image dimensions and Z-section thickness. The resulting images had X and Y pixel sizes in the range of 30-40nm and Z sizes in the range of 140-500nm depending on lens/zoom combinations. Images were processed using Imaris software v10.1 (Oxford Instruments, Boston, MA), employing the 3D-visualization option along with surface segmentation, classifier sub-segmentation routines, and slicers as required. Image intensity was corrected in Imaris, and image files were exported as TIFF files. Composite images were prepared in Adobe Illustrator CC2024. Videos were generated using the Animation tools provided in the Imaris software package. Images are representative of experiments performed in 5-10 rats.

### Quantitation of desmosome parameters in filled and voided bladders

The Navigator function of Leica LASX software, in conjunction with the 10X objective (zoom set to 1.0), were employed to generate images of entire filled and voided rat bladders immunolabeled with antibodies to DSG2 and stained with phalloidin as a marker of the AJC-associated actin ring (and lateral cell surface). The Navigator tool was used to superimpose a grid on the image (each square encompassing an area of ∼ 1165 x 1165µm) and a random number table was used to identify 10 areas for further analysis. Images obtained from the center of each of these 10 areas were collected (63X objective, zoom=1, ∼ 185 x 185 µm area) using the LIGHTNING deconvolution module described above. If multiple whole cells were present in an image, a random number table was used to identify cells for further analysis. Z-stacks of final images were imported into Imaris, and the Surfaces tool of the 3D visualization module was used to segment the DSG2-labeled desmosomes. The Classification option was then used to select the AJC-associated desmosomes. The following parameters were collected for classified desmosomes: number, area, and volume. In addition, the Imaris Filament tool was used in conjunction with the AJC-associated actin and DSG2 labeling to define and quantify the cell perimeter. Values for each animal were averaged and the mean and SEM of the composite data reported.

### PREM and SEM

PREM was performed using a previously described protocol by Svitkina ^46^, but with the following modifications. The original protocol calls for the use of Triton X-100 to remove membranes, but the umbrella cell apical membrane is highly detergent resistant and is not solubilized by Triton-X-100 ^76^. We also found that octylglucoside and digitonin were also ineffective. In contrast, the non-ionic detergent POE effectively removed membrane while preserving the cytoskeleton. An additional, key modification to the published PREM protocol included reducing the concentration of polyethylene glycol, a cytoskeleton stabilizing agent, in the extraction solution from 2.0% to 0.2%. At the higher concentration, a large amount of residual protein adhered to the cytoskeleton, obscuring its visualization. The lower concentration effectively maintained the cytoskeletal architecture while eliminating much (but not all) of the adherent protein.

In our studies, we used unfixed and stretched bladder tissue, prepared as described above. The tissue was rinsed with PBS and then treated with freshly prepared PCS buffer supplemented with 2µM unlabeled phalloidin for 45 min at room temperature on a slowly rotating orbital shaker (30rpm). The tissue was rinsed with PEM buffer three times, rotating the dish by hand after each wash. The sample was then treated for 30min at room temperature with 1mg/ml S1 fragments derived from rabbit psoas skeletal muscle (Cytoskeleton Incorporated, Denver, CO; CS-MYS04), reconstituted in PEM buffer containing 2.0µM phalloidin and centrifuged at 100,000 x g for 15min at 4°C prior to use. The tissue was then rinsed with PEM buffer, then PBS, and then fixed in 2% v/v glutaraldehyde in phosphate buffer for 30min at room temperature. The tissue was then rinsed with distilled water, treated with 0.1% w/v tannic acid (EMS) for 20min, rinsed twice with distilled H2O, washed with distilled H2O for 5min, treated with 0.2% w/v aqueous uranyl acetate for 20min at room temperature, and then dehydrated using ethanol. The sample was then treated with 0.2% w/v uranyl acetate (prepared in ethanol) for 20min at room temperature, and then dehydrated further using 100% ethanol for 5min, followed by two 5-min incubations with molecular-sieve-dried ethanol (prepared as described ^46^). The samples were then critical point dried using a Samdri-PVT-3D critical point dryer (Tousimis, Rockville, MD) fed with bone-dry CO2. For PREM, the dried samples were transferred to a JEOL JFD-V-TP device, rotary shadowed with ∼ 2nm of platinum at 45°, followed by shadowing with ∼ 5.0nm of carbon. As household bleach was often ineffective at removing residual tissues from replicas, they were instead treated with a 1:10 dilution of sodium hypochlorite (4.0-4.99% available chloride) overnight at room temperature in porcelain spot plates to remove remaining tissue. The replicas were washed with H2O three times and then picked up on Butvar-coated nickel grids and viewed in a JEOL1400 FLASH TEM. For SEM, the critical point dried samples (taken prior to rotary shadowing) were sputter coated with gold palladium using a Cressington 108auto sputter coater (Ted Pella, Redding, CA), mounted on aluminum stubs, and viewed in a JEOL 6335F field-emission SEM. Immunogold labeling was performed as described by Svitkina et al ^46^. The PREM and SEM images presented are representative of experiments performed on 5-10 rats.

### Treatment with CytoD, LatA, nocodazole, and plecstatin-1

Stretched bladder tissue was prepared as described above. After the 30-min equilibration in Krebs buffer, the tissue was treated for 60min at 37°C with 33µM nocodazole, 25µg/ml CytoD, 1-50µM LatA (Biotechne Tocris Bioscience, Minneapolis, MN), 25µM plecstatin-1-HCl (MedChemExpress, Monmouth Junction, NJ), or DMSO dissolved in Krebs buffer. All drugs were prepared as 1000-fold stock solutions, stored as aliquots at -20° C, and diluted just prior to use.

### Detection of PLEC variants in rodent umbrella cells using PCR

For mouse urothelium, the following protocol was employed. Mouse bladders were excised and rinsed with Kreb’s buffer. The pointed end of a yellow tip, trimmed by 5mm with a scalpel, was positioned next to the dome of the bladder. Fine forceps were used to invert the bladder onto the tip (mucosal surface now facing out). The inverted bladder was placed in 150µl of lysis/binding buffer (RNAqueous-4PCR Kit; Invitrogen, Waltham, MA) for 30s and total RNA was isolated per the manufacturer’s protocol. An AccuSript PfuUltra II RT-PCR kit (Agilent, Santa Clara, CA) was used to generate cDNAs using random primers and following the manufacturer’s protocol. The splice variant-specific primers employed for PCR analysis were those provided by Rezniczek *et al* ^39^. PCR reactions, were performed using the KAPA HiFi polymerase kit (Roche Sequencing Solutions, Pleasanton, CA) and a T100 Thermal Cycler (BioRad, Hercules, CA) using the following protocol: initial denaturation at 94°C for 2min, followed by 39 cycles of 94°C for 20s, 60°C for 15s, 72°C for 15s, and a final step of 72°C for 2min prior to holding at 4°C. Amplicons were resolved using 2% w/v agarose gels.

For rat umbrella cells, the following protocol was performed. Stretched rat bladders were pinned out on rubber mats and treated with 2.5mg/ml of Dispase (Gibco-ThermoFisher Scientific) dissolved in Minimal Essential Medium (Gibco-ThermoFisher Scientific) for 60 min at 37° C in a cell culture incubator. The mucosal surface was gently scraped using blunt forceps, and the medium with cells collected and centrifuged at 200 x g in a table-top centrifuge for 5min. The cell pellet was resuspended in Kreb’s buffer, and while viewing through a microscope an oil-filled glass pipette (prepared from borosilicate glass capillary tubes using a PP-81 puller; Narishige International USA, Amityville, NY), attached to a Nanoject II suction system (Drummond Scientific, Broomall, PA), was used to collect a total of 30 umbrella cells. The latter were identified by their large size and binucleate morphology. Cell lysis and cDNA synthesis was performed using the SuperScript IV CellsDirect cDNA Synthesis kit (ThermoFisher Scientific). PCR was performed as described above, but using the following primers:

> RAT-Plec1-Fwr: TGACCTCGCTGAAAGCTCG
>
> RAT-Plec1-Rev: GGTCTCGTTCATCTGTGGCT
>
> RAT-Plec1a-Fwr: GGTAGCAAGAGAACCAGCTCA
>
> RAT-Plec1a-Rev: AGGTGTTTATTGACCCACTTG
>
> RAT-Plec1c-Fwr: GAGTGGAGGTGGTTCTGTGG
>
> RAT-Plec1c-Rev: AGGTGTTTATTGACCCACTTG
>
> RAT-Plec1f-Fwr: CCGACGAACAGGACTTCATC
>
> RAT-Plec1f-Rev: AGGTGTTTATTGACCCACTTG

### Generation of PLEC1-8-EGFP and PLEC1a6-8-EGFP adenoviruses and *in-situ* adenoviral transduction

Gene synthesis, subcloning into adenoviral expression vectors, and adenovirus production of the PLEC1a-8-EGFP and PLEC1a6-8-EGFP constructs was performed by VectorBuilder (Chicago, IL). For the PLEC1a-8-EGFP construct, a cDNA was synthesized that encoded amino acids 1-275 of the rat PLEC1a variant (transcript variant 11, NCBI Reference Sequence: NM_001164308.3; exons 1-8), followed by the in-frame addition of a 5’-ggcagcgcgggcagcgcggcgggcagcggcgaattt sequence (encoding a GSAGSAAGSGEF flexible linker) and the cDNA sequence encoding EGFP. The PLEC1a6-8-EGFP construct was identical but lacked exons 2-5 (which encode amino acids 38-151), generating a construct that lacked the calponin-homology domain1 actin binding motif of PLEC1a. The synthesized constructs, with included Kozak sequences, were cloned into the pAVexpression vector behind a CMV promoter and adenoviruses were produced using the pilot-scale packaging service. Upon receipt, the adenoviruses were further amplified using our previously described methods and stored at -80° C prior to use ^77^.

*In situ* transduction was performed as described previously ^56, 77^. Briefly, mice were sedated with 2.5% (v/v) isoflurane and a 22-g Jelco IV catheter, with Luer fitting (Smith Medicals) and trimmed to ∼ 1-cm in length, was inserted into the bladder via the urethra. The bladder was rinsed with 100µl of PBS delivered via a 1-ml syringe, which was then detached to allow the buffer to void. The bladder was then filled with 100μl of 0.1% w/v dodecyl-β-D-maltoside dissolved in PBS via syringe. The syringe was left attached to the catheter for 10min to prevent the solution from escaping. The bladder was allowed to void, and then was filled with 100μl PBS via syringe containing adenoviruses expressing the constructs described above (1.0 x 10^8^ infectious virus particles per animal). The syringe was left in place, and after 30min detached and the virus solution was allowed to void. Anesthesia was discontinued, and the mice were allowed to revive. At 48-h post transduction, the mice were subjected to the filled bladder protocol described above with the following changes. Mice were anesthetized with 2.5% v/v isoflurane and the mouse bladders were filled to 150µl over 30 min prior to perfusion fixation. Samples were processed for immunofluorescence as described above.

## Statistical analysis

Data are reported as mean ± SEM. Statistically significant differences were determined using two-tailed Student t-tests. A p value ≤ 0.05 was considered statistically significant.

## ACKNOWLEDGEMENTS

This work was supported by grants from the National institutes of Health, including R01DK129473 (to GA) and by the Pittsburgh Center for Kidney Research KIDNIT imaging core (U54DK137329). The Leica Stellaris confocal used in this study was funded in large part by S10OD028596 (to GA).

## AUTHOR CONTRIBUTIONS

Conceived and designed research: WR, DC, MD, GA

Performed experiments, analyzed data, and interpreted results of experiments: WR, DC, TP, MD, JF, GA

Prepared figures: DC, MD, GA Drafted manuscript: GA, MD

Edited and revised manuscript: GA, MD, DC

Approved final version of manuscript: WR, DC, TP, MD, JF, GA

