## Supplementary Text and Figures for "The umbrella cell keratin network: organization as a tile-like mesh, formation of a girded layer in response to bladder filling, and dependence on the plectin cytolinker"

#### SUPPLEMENTARY FIGURE LEGENDS

**Fig. S1.** Distribution of KRT8 and KRT20 in umbrella cell. **(A-B)** Images of tricellular region of umbrella cells obtained from filled bladders acquired using confocal-SRIP microscopy coupled with 3D reconstruction of the keratin network. Panel A is a 3D-reconstruction of a tricellular region and panel B shows individual X-Y optical sections at the indicated position in Z.

**Fig. S2.** Distribution of KRT20 in an umbrella cell taken from a bladder filled to near maximal capacity (1000 µl). Upper image is a 3D reconstruction of a Z-series collected using confocal-SRIP microscopy. Smaller images below are individual XY sections taken at the indicated Z depth (top of cells set to Z=0). The positions of the umbrella cell nuclei are indicated by white circles. Other nuclei are those of intermediate cells.

**Fig. S3.** Localization of ACTN4 and KRT8 in bladder umbrella cell. **(A-C)** Image of umbrella cell from a filled bladder that was acquired using confocal-SRIP and then reconstructed in 3D. **(A)** The smaller images to the right show data for individual markers. Boxed regions are reproduced in panel B. **(B)** The boxed regions are magnified in the images below. **(C)** Arrowheads point to the location of the AJC-associated actin ring, which is $ACTN4^{+}$ but $KRT8^{-}$.

**Fig. S4.** Effects of CytoD, LatA, and nocodazole on the umbrella cell keratin network. Stretched bladder tissue was treated with DMSO (solvent control), 25 µg/ml CytoD, 25 µM LatA, or 33 µM nocodazole (Noc) for 60 min. Images were acquired using confocal microscopy and 3D reconstructions generated. Boxed regions are magnified in the insets. Capturing the complete outlines of the umbrella cells, which are often in different Z planes, necessitated including the underyling intermediate cell layer.

**Fig. S5.** Association between microtubules and PLEC in umbrella cells. **(A)** Tricellular region of umbrella cells from a filled bladder showing the dispersion of microtubules within the apical keratin network. Image was acquired using confocal-SRIP imaging and a 3D reconstruction with rendered surfaces was generated. Only the PLEC associated with the AJC is displayed. **(B)** Same region as shown in panel A; however, without surface rendering and with all PLEC signal displayed. The boxed region is magnified in the inset. Note the close association of PLEC with microtubules. **(C)** Same view as A, but without the KRT20 signal. Circled areas: regions of close contact between microtubules and the AJC. Boxed region is magnified in inset.

**Fig. S6.** *Plec* splice variant expression in rat umbrella cells. RNA was purified from isolated umbrella cells and the indicated splice variants of *Plec* amplified using RT-PCR.

**Fig. S7.** Effect of expressing PLEC1a8-EGFP or PLEC1a6-8-EGFP on actin and PLEC expression and distribution. Mouse bladders where transduced with viruses encoding the indicated construct. Images were acquired using confocal imaging and 3D reconstruction was performed. Note the presence of visible actin filaments in the bottom left cells expressing larger amounts of PLEC1a8-EGFP. AJC-associated PLEC is lost in cells expressing PLEC1a8-EGFP. The construct PLEC1a-6-8-EGFP does not appear to affect the expression or distribution of actin or PLEC.

**Fig. S8.** Effects of plecstatin-1 treatment on umbrella cell AJC continuity and morphology. **(A)** Stretched bladders were treated with DMSO solvent or 25 µM plecstatin-1 for 60 min prior to processing for confocal microscopy and 3D rendering. Yellow arrows point to regions of the AJC exhibiting discontinuity of the CLDN8 and actin rings. White arrowheads point to the location of a mesa-like focus. Dashed region in the upper right of the “merge” image is missing umbrella cells, which have detached, exposing the underlying intermediate cell layer. **(B-C)** PREM imaging of plecstatin-1-treated umbrella cell layer. **(B)** Formation of mesa-like foci (marked with white asterisks) in plecstatin-1-treated umbrella cells. The upper mesa is nearer the center of cell "1," while the lower mesa is near its AJC border. Arrow indicates the loss of cell-cell cohesion between cells "3" and "4." The boxed region is magnified in panel C. **(C)** Near the mesa, the apical cytoskeleton of plecstatin-1-treated cells contains abundant crisscrossed/woven S1-decorated actin filaments along with interspersed intermediate filaments and cytolinkers.

#### SUPPLEMENTARY VIDEOS:

**Video1:** 3D rendering of umbrella cell AJC at a tricellular region. TJP1, green; actin, red; CDH1, white; AJC-associated desmosomes, magenta; lateral desmosomes cyan; basal desmosomes dark blue.

**Video2:** Apical keratin network of umbrella cell taken from filled bladder.

**Video3:** Girded layer of keratin network taken from umbrella cell of bladder filled to 1000µl.

**Video4:** Keratin network of umbrella cell taken from quiescent bladder.

**Video5:** Relationship of KRT20 network to DSP/DSG2-labeled desmosomes.

**Video6:** Distribution of PLEC in umbrella cells. AJC-associated PLEC, green; PLEC away from AJC, dark blue; actin, red; KRT20, white.

#### Figure S1

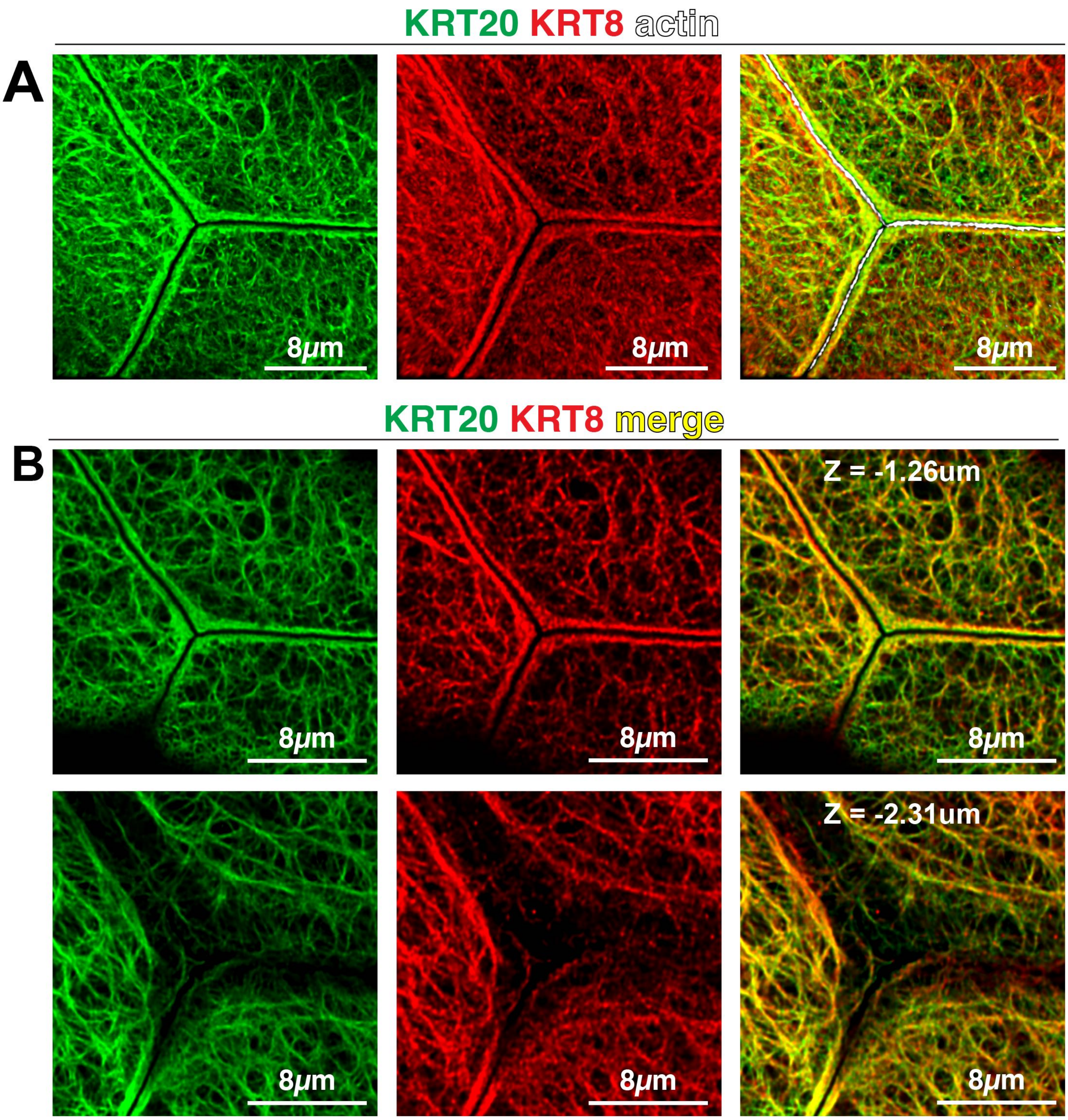

KRT20 KRT8 actin
A
8μm
8μm
8μm
KRT20 KRT8 merge
B
Z = -1.26um
8μm
8μm
8μm
Z = -2.31um
8μm
8μm
8μm

Figure S2

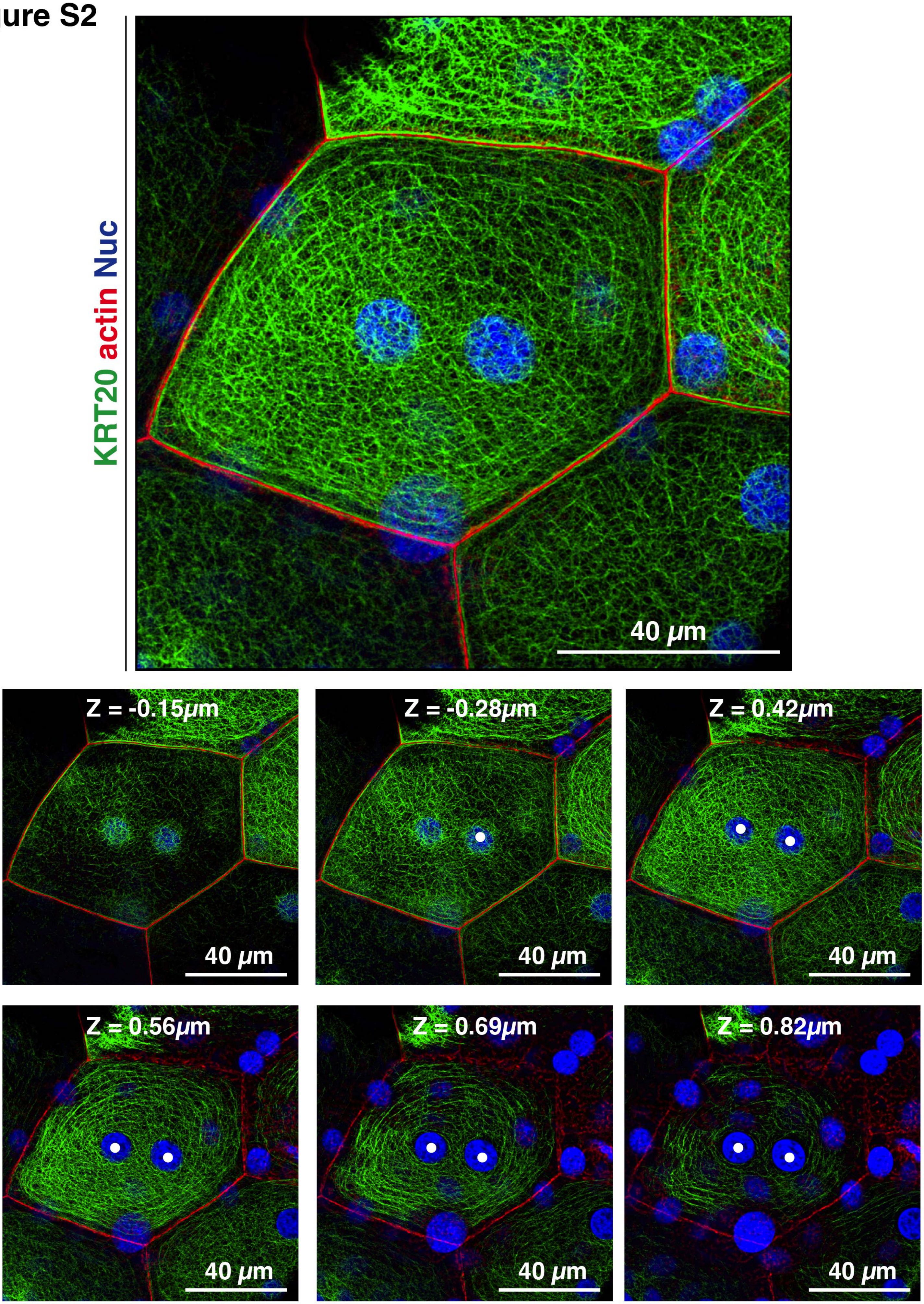
KRT20 actin Nuc
40 μm
Z = -0.15μm
40 μm
Z = -0.28μm
40 μm
Z = 0.42μm
40 μm
Z = 0.56μm
40 μm
Z = 0.69μm
40 μm
Z = 0.82μm
40 μm

Figure S3

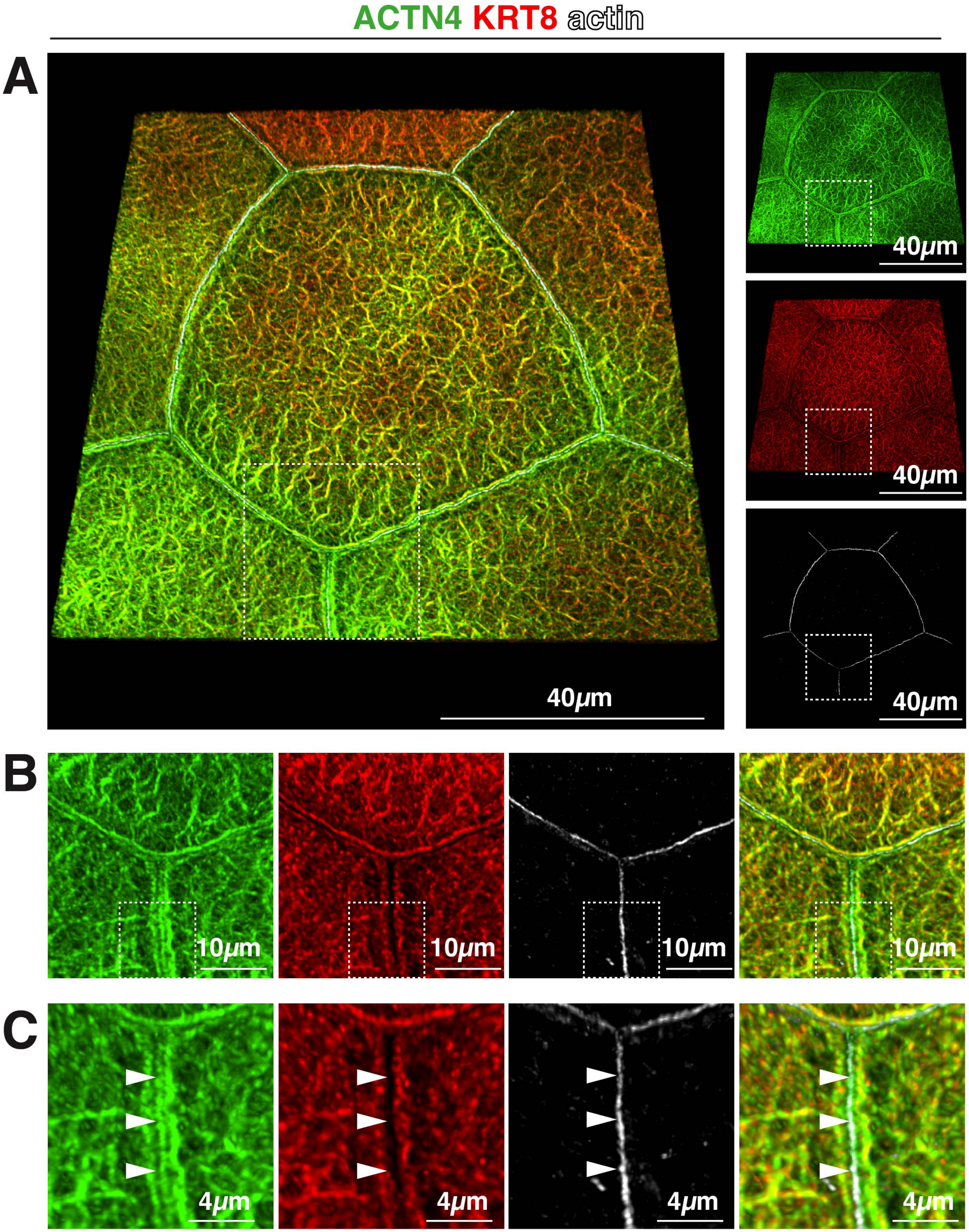

ACTN4 KRT8 actin
A
40μm
40μm
40μm
40μm
B
10μm
10μm
10μm
10μm
C
4μm
4μm
4μm
4μm

### Figure S4

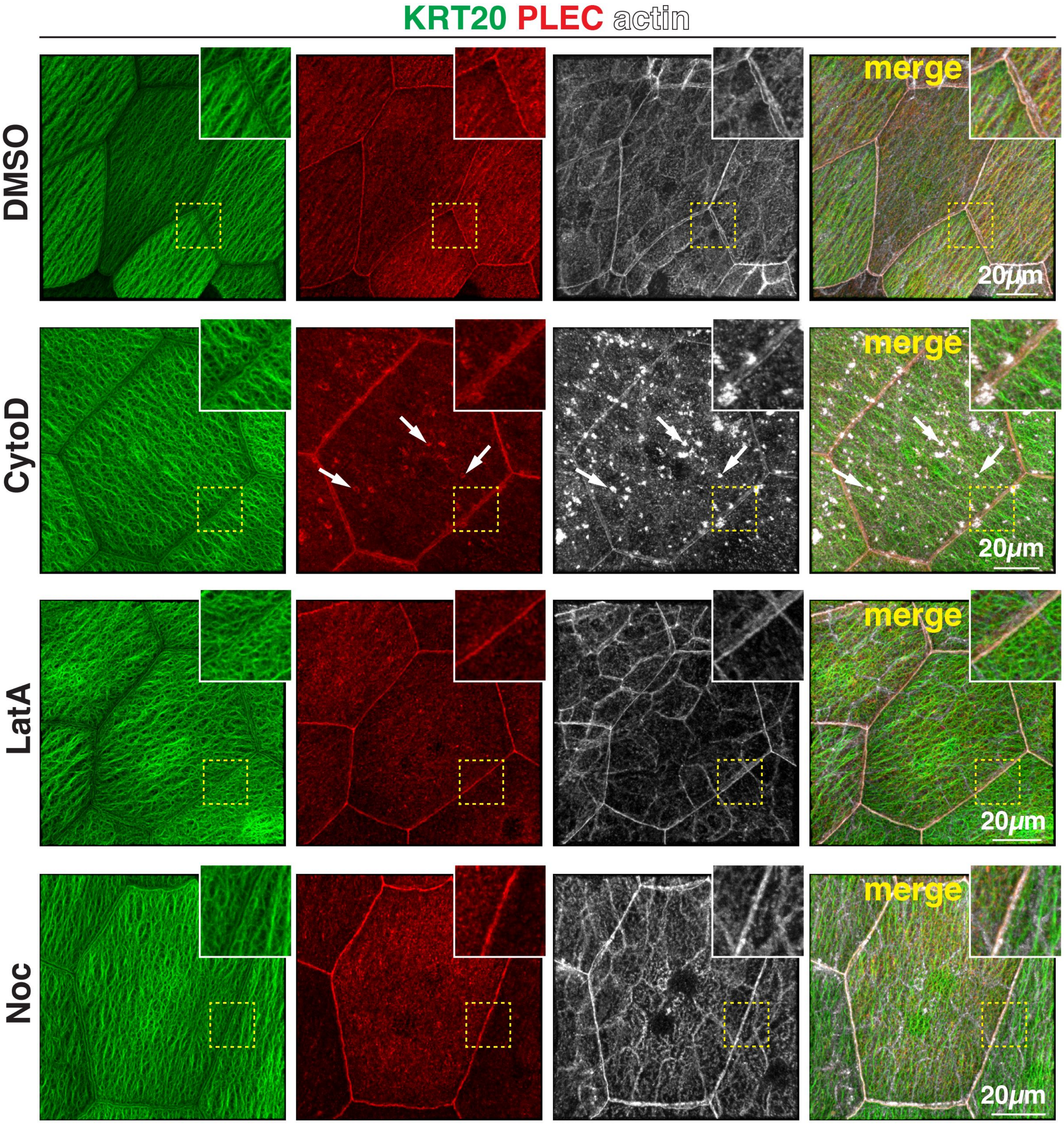
KRT20 PLEC actin
DMSO
CytoD
LatA
Noc
merge
merge
merge
merge
20μm
20μm
20μm
20μm

**Figure S5**

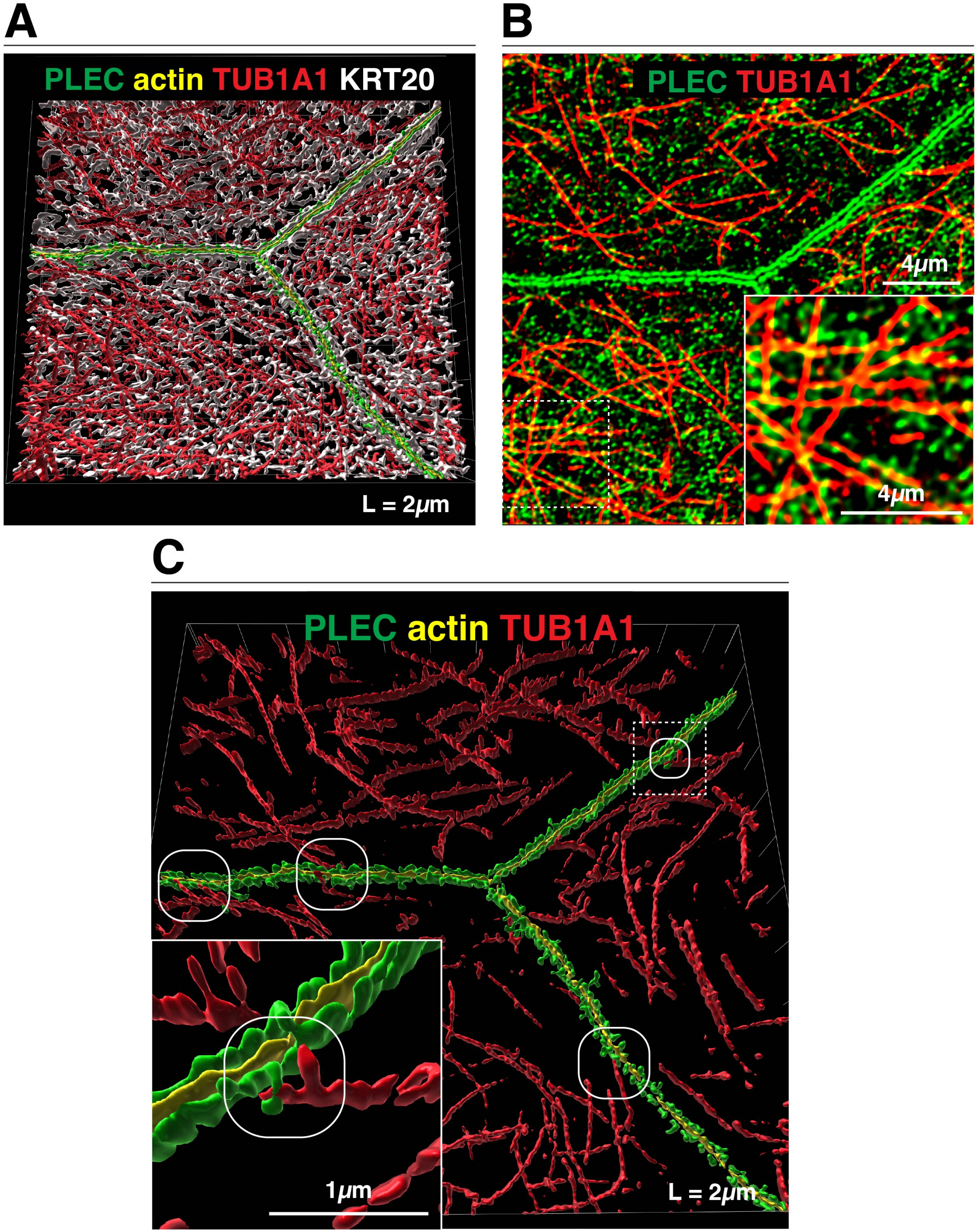
A
PLEC actin TUB1A1 KRT20
L = 2μm
B
PLEC TUB1A1
4μm
4μm
C
PLEC actin TUB1A1
1μm
L = 2μm

### Figure S6

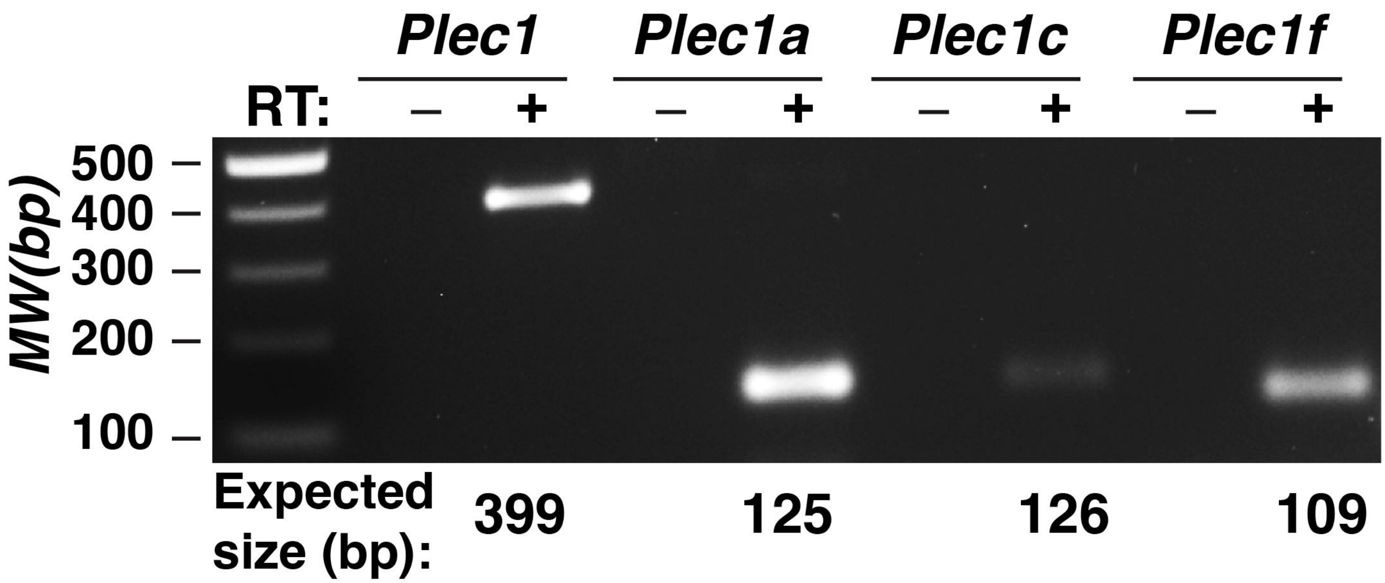

Plec1
Plec1a
Plec1c
Plec1f
RT:
–
+
–
+
–
+
–
+
500
400
300
200
100
MW(bp)
Expected size (bp):
399
125
126
109

**Figure S7**

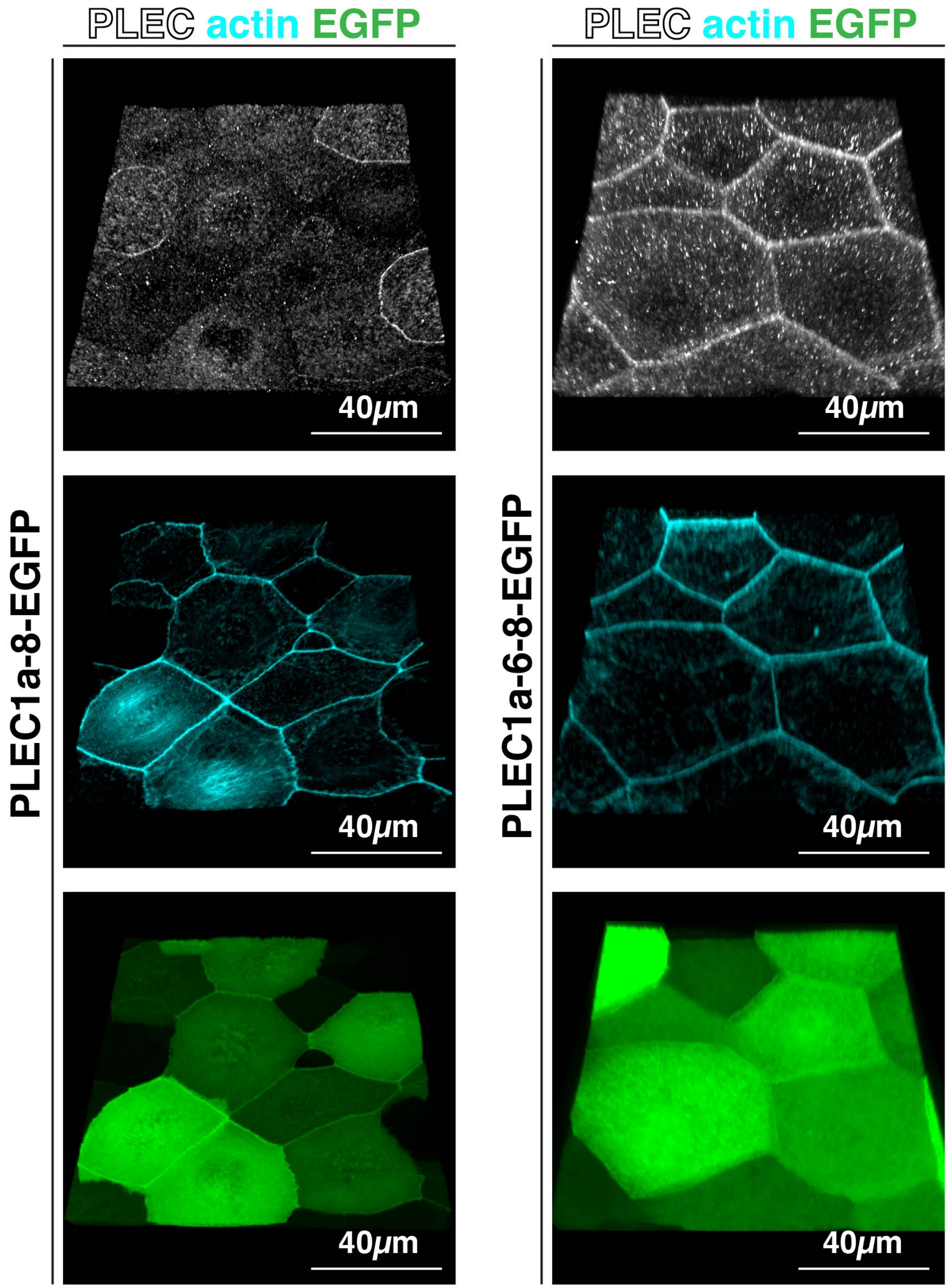
PLEC actin EGFP
PLEC actin EGFP
PLEC1a-8-EGFP
PLEC1a-6-8-EGFP
40μm
40μm
40μm
40μm
40μm
40μm

**Figure S8**

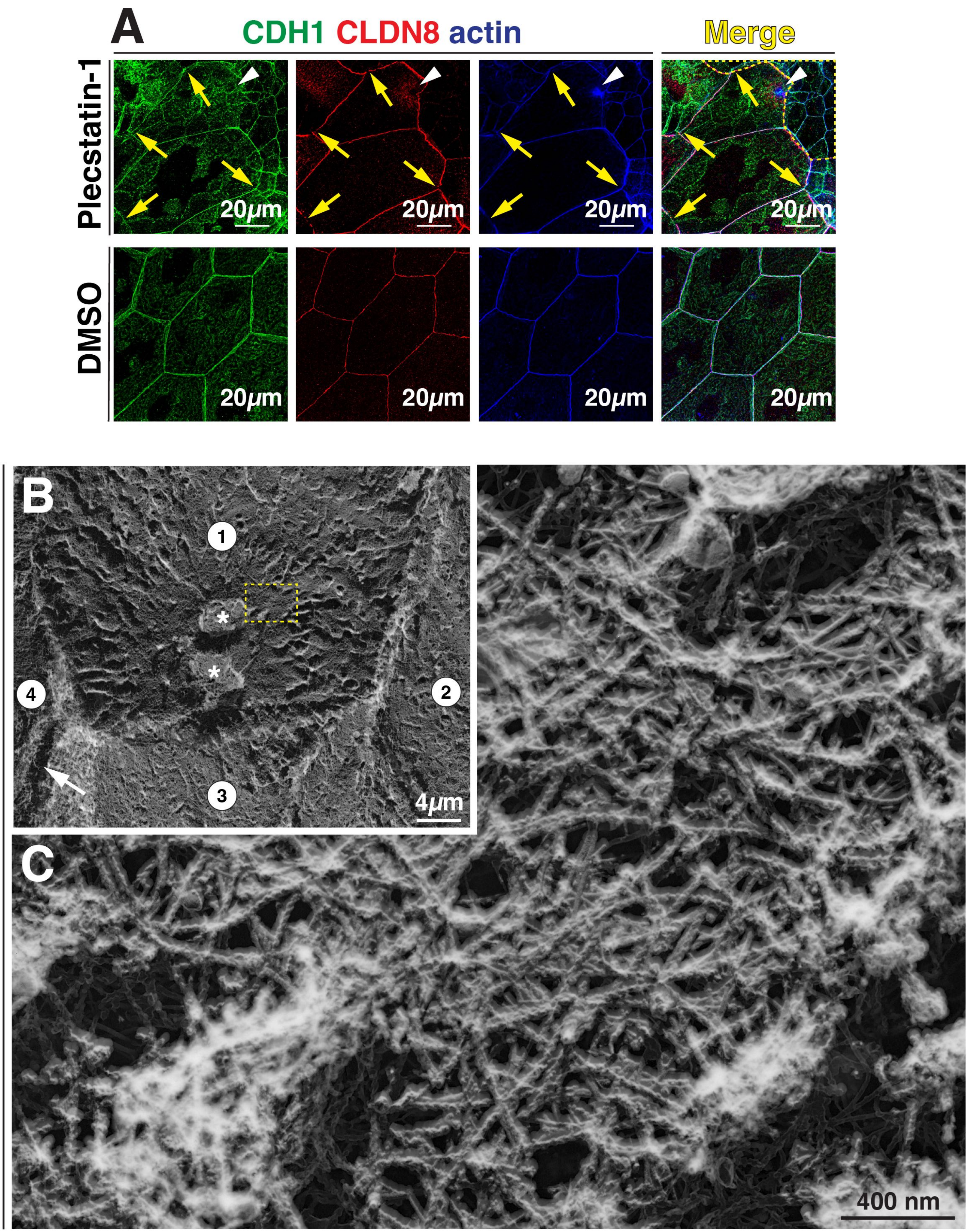

A
CDH1 CLDN8 actin
Merge
Plecstatin-1
DMSO
20μm
B
1
2
3
4
*
4μm
C
400 nm
